# Engineered collagen-targeting therapeutics reverse lung and kidney fibrosis in mice

**DOI:** 10.1101/2022.01.04.474747

**Authors:** Michael JV White, Michal M Raczy, Erica Budina, Eiji Yuba, Ani Solanki, Ha-Na Shim, Zheng Jenny Zhang, Laura T Gray, Shijie Cao, Aaron T. Alpar, Jeffrey A Hubbell

## Abstract

Fibrotic diseases are involved in 45% of deaths in the United States. In particular, fibrosis of the kidney and lung are major public health concerns due to their high prevalence and lack of existing treatment options. Here, we harness the pathophysiological features of fibrotic diseases, namely leaky vasculature and aberrant extracellular matrix (ECM) protein deposition (i.e. collagen), to target an anti-fibrotic biologic and a small molecule drug to disease sites of fibrosis, thus improving their therapeutic potential in mouse models of lung and kidney fibrosis. First, we identify and validate collagen-targeting drug delivery systems that preferentially accumulate in the diseased organs: von Willebrand Factor’s A3 domain (VWF-A3) and decorin-derived collagen-binding peptide-conjugated micelles (CBP-micelles). We then engineer and recombinantly express novel candidate biologic therapies based on the anti-inflammatory cytokine IL-10: A3-IL-10 and A3-Serum Albumin-IL-10 (A3-SA-IL-10). Simultaneously, we stably encapsulate the potential anti-fibrotic water-insoluble drug, rapamycin, in CBP-micelles. We show that these novel formulations of therapeutics bind to collagen *in vitro* and that their efficacy in mouse models of lung and kidney fibrosis is improved, compared to free, untargeted drugs. Our results demonstrate that collagen-targeted anti-fibrotic drugs may be next generation therapies of high clinical potential.

## 1. Introduction

Fibrosing diseases—including pulmonary fibrosis, congestive heart failure, liver cirrhosis, and end-stage kidney disease—are involved in 45% of deaths in the United States [1, 2]. Of these diseases, kidney and lung fibrosis, caused by the excessive production and deposition of extracellular matrix (ECM) proteins in the kidney and lung interstitium, respectively, are among the most challenging to treat and eventually lead to end-stage kidney disease (ESKD) and lung failure. The two FDA approved treatments for fibrosis, pirfenidone and nintedanib, [2, 3] slow, but fail to reverse, the progression of fibrosis via poorly understood mechanisms of action [4]. Currently, the only intervention capable of restoring kidney and lung function is organ transplantation [5]. However, the number of patients awaiting a transplant far exceeds the available supply of donor organs [5]. Hence, there remains an unmet need for new therapies that can better target and treat the fibrotic organs.

Phenotypically, fibrosis occurs when tissue injury activates macrophages to secrete growth factors, enzymes, and cytokines that stimulate fibroblast migration and proliferation [6]. Recruited fibroblasts, in turn, produce large amounts of extracellular matrix (ECM) proteins including collagens type I and III and fibronectin, resulting in tissue that is rich in collagen and characterized by leaky vasculature [8-11]. Kidney and lung fibrosis progression can be mirrored in mice through several injury models, including the unilateral ureteral obstruction (UUO) model, in which one of the kidneys is blocked via ligation of the ureter [12-14], and the bleomycin-induced pulmonary fibrosis model, in which the interstitial edema of the lungs is compromised by the agent bleomycin, respectively [15].

A number of protein and small molecule-based therapies are currently being explored for the treatment of fibrosing diseases. Interleukin-10 (IL-10) is an immunosuppressive cytokine that plays a critical role in preventing inflammatory and autoimmune diseases [16]. When IL-10 binds to the heterodimeric IL-10 receptor, the JAK/STAT signaling pathway is activated, suppressing the production of pro-inflammatory mediators including TNF-*α* and IL-1*β* [16]. IL-10 treatment also reportedly reduces antigen presentation and phagocytosis by antigen presenting cells and inhibits pathogenic T helper cell proliferation, while expanding immunosuppressive regulatory T cells [7]. Due to its established role in suppressing macrophage and myofibroblast-mediated inflammation, IL-10 is a promising candidate for cytokine therapy in fibrosis [18]. IL-10 gene and recombinant protein delivery has been reported to suppress bleomycin-induced pulmonary fibrosis by inhibiting the production of TGF-β1 by alveolar macrophages, lung fibroblasts, and myofibroblasts [8, 9]. Treatment with recombinant IL-10 has also been reported to reduce liver fibrosis in chronic hepatitis C patients who did not respond to interferon-based therapy [10]. Despite these promising results, the therapeutic utility of recombinant IL-10 has been hindered by the inability to reach sufficiently high local concentrations within the fibrotic microenvironment.

Rapamycin (sirolimus) is a natural macrocyclic lactone used clinically as an orally-administered immune suppressant that operates through inhibition of the mammalian target-of-rapamycin (mTOR) receptor [11, 12]. Rapamycin is an anti-inflammatory and autophagy-inducing molecule, and has been reported to inhibit the progression of both kidney [13] and lung [14] fibrosis via increased expression of TGF-α and epidermal growth factor receptor (EGFR) signaling. However, the therapeutic use of intravenously infused rapamycin is hindered by its low solubility in water and the difficulty of its stable formulation, resulting in unwanted systemic side effects [12].

Here, we engineer delivery systems that target sites of exposed collagen prevalent in kidney and lung fibrosis. To target the cytokines to the leaky vasculature of fibrotic tissues, we used protein engineering to recombinantly fuse the von Willebrand Factor A3 domain (A3-VWF), a collagen-binding domain, to the cytokine IL-10, and also engineered collagen binding peptide-conjugated micelles for the encapsulation of the small molecule rapamycin. We further compared collagen-targeting with targeting a splice variant of fibronectin (FN), the fibronectin Extra Domain A (FN-EDA) domain. We then evaluated the targeting ability of these drug delivery systems and demonstrated superior efficacy to unmodified counterparts in mouse models of fibrosis. These findings support the use of innovative collagen-targeting approaches to improve the clinical potential of anti-fibrotic drugs.

## 2. Results and Discussion

The leakiness of the fibrotic tissue vasculature exposes ECM proteins normally unavailable in blood circulation. In this work, we harnessed this phenomenon to actively target therapies to sites of fibrosis. We sought to target IL-10, an anti-inflammatory cytokine, or rapamycin, a small molecule anti-fibrotic. Our laboratory has previously identified several peptide and protein-based ligands that bind to exposed ECM and target to tissues: 1) von Willebrand Factor A3 domain (VWF-A3), which binds to collagens I and III [15]; 2) a fibronectin-extra-domain-A (FN-EDA) antigen-binding fragment (α-FN-EDA Fab) with affinity for FN-EDA [28, 29]; and 3) decorin-derived collagen-binding peptide (CBP) that binds to collagens I and III [16]. While the protein IL-10 can be fused to VWF-A3 or α-FN-EDA Fab targeting domains via recombinant protein engineering, the small molecule rapamycin is water insoluble and requires additional encapsulation. For this purpose, we developed a CBP-conjugated micelle for the stable encapsulation of rapamycin.

### 2.1. Rapamycin-CBP-micelle Development and Characterization

Rapamycin is a well-validated anti-inflammatory small-molecule antibiotic that is being explored as an anti-fibrotic drug due to its ability to inhibit fibroblast proliferation [32, 33]. However, its low water solubility makes the formulation process difficult and its low bioavailability results in the systemic side effects [24]. As a result, the use of rapamycin as intravenously-administered drug is limited. We developed a targeted drug delivery system engineered to both efficiently solubilize rapamycin and target it to the fibrotic, collagen-rich tissues. To encapsulate rapamycin, we use a block copolymer of poly(ethylene glycol) (pEG) and poly(phenylalanine) (pPhe) which forms nanometer-scale micelles that have been reported good carriers of hydrophobic small molecule drugs [34]. The pEG_114_-pPhe_20_ polymer of about 8kDa was synthesized as previously described [34] (Supplementary Figure 1A), and its block lengths confirmed by 1HNMR (Supplementary Figure 1B) and MALDI-TOF (Supplementary Figure 1C). The polymer forms micelles when dispersed in water at the concentration above their critical micellar concentration (CMC) of 0.01mM (Supplementary Figure 2A). Hydrophobic compounds encapsulated in pEG_114_-pPhe_20_ micelles are released through passive diffusion following Korsmeyer-Peppas drug release kinetic, as demonstrated by *in vitro* release kinetic study using model small molecule dye rhodamine B (Supplementary Figure 2B).

In our approach, we additionally linked the micelles with a decorin-derived collagen binding peptide (CBP, CNNNLHLERL) via a small-molecule heterobifunctional linker (Supplementary Figure 3). First, peptide was reacted with the linker to arm it with azide-reactive dibenzocyclooctine (DBCO) functional group, yielding peptide-DBCO. Next, azide-modified micelles were reacted with a peptide-DBCO, resulting in peptide-micelle conjugate. As a non-binding control, a scrambled-sequence peptide (SCR, CLLNEHLNNR) was conjugated to the micelles in place of CBP. Peptide-micelle conjugates were first characterized by Dynamic Light Scattering (DLS) to determine nanoparticle size (Supplementary Figure 4A). We observed an increase in micelle diameter from 18.0 nm to 24.6 nm and 24.6 nm upon CBP and SCR peptide conjugation, respectively (Supplementary Figure 4A). In addition, cryogenic electron microscopy (cryo-EM) images of the conjugates confirmed their size of about 15-20 nm and demonstrated their semi-spherical shape (Supplementary Figure 4B). There were no obvious differences observed between cryo-EM images of conjugated and unmodified micelles (Supplementary Figure 4B). The presence of CBP peptide conjugated to nanoparticles was evidenced by performing a reactive site-competition reaction experiment. First, azide-modified micelles were either reacted with peptide-DBCO or left unreacted. Subsequently, azide-reactive fluorescent dye (AF647-DBCO) was added to both peptide-reacted and unreacted micelles; reaction could happen only if free azide groups were available. Based on size exclusion chromatography (SEC) elution profiles (Supplementary Figure 5A) we observed that AF647-DBCO reacted only with unreacted micelles (fractions 15-25), providing a proof of a successful conjugation of the peptide. Next, we examined if the CBP micelles could bind to collagen *in vitro* to enable their accumulation in collagen-rich environment (Supplementary Figure 5B). To demonstrate this, we incubated fluorescently labeled CBP-micelles or SCR-micelles (non-binding control) with collagen type I sponge hydrogels (Helistat, cut-out to ca. 0.2 cm in diameter). After an hour of incubation at 37°C, the gels were taken out, washed with PBS and imaged using IVIS imaging. Collagen gels incubated with CBP-micelles had significantly higher fluorescence than gels incubated with the SCR-micelles, which demonstrated the presence of the peptide on the micelles and the preserved affinity to collagen type I (Supplementary Figure 5B).

### 2.2. VWF-A3 and CBP-micelles Target to Both Fibrotic Kidney and Lungs

Next, we tested the ability of these novel peptide and antibody targeting agents to target to either the fibrotic kidney, induced by unilateral ureteral obstruction (UUO) or the fibrotic lung, induced by bleomycin instillation (Figure 1). For this purpose, we insulted mouse kidneys or mouse lungs in two separate studies and allowed fibrosis to develop in these organs for 1 week. We then intravenously injected fluorescently labeled (1) α-FN-EDA Fab, (2) VWF-A3, (3) CBP-micelles and (4) scrambled CBP-micelles at equimolar concentrations and compared the fluorescence of the harvested organs after 24 hours or 48 hours for kidney fibrosis or lung fibrosis respectively via IVIS (Figure 1, Supplementary Figures 6, 7).

**Figure 1.**
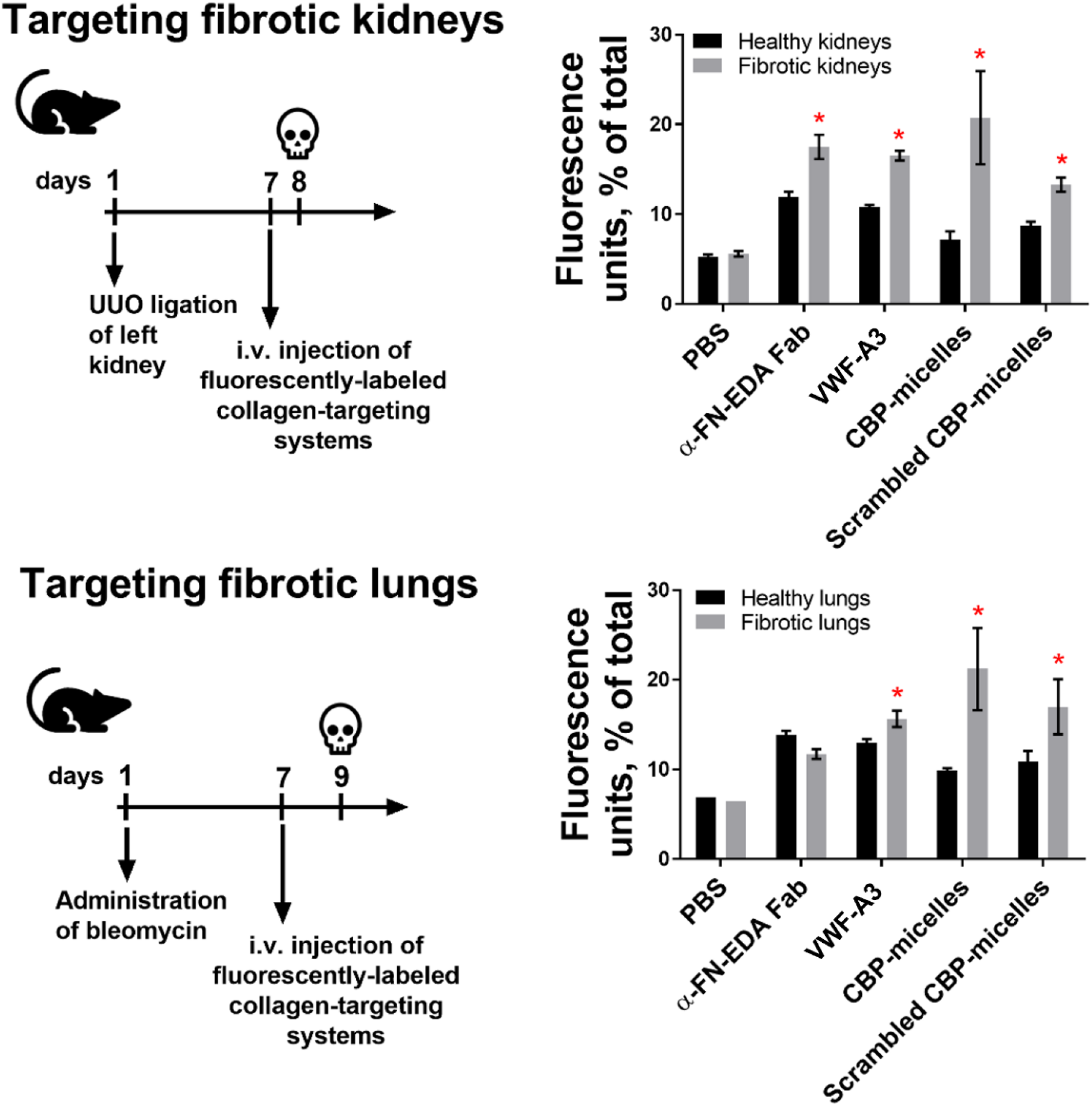
Fluorescently labeled VWF-A3 and CBP-micelles target to both fibrotic kidney and lungs. The left kidneys of mice were insulted with UUO ligation, and the lungs of other mice were insulted with intranasally instilled bleomycin (75 μg). These mice were allowed to become fibrotic for 1 week and were then injected i.v. with fluorescently labeled (1) α-FN-EDA Fab, (2) VWF-A3, (3) CBP-micelles or (4) Scrambled CBP-micelles at equimolar concentrations. Heart, lung, spleen, kidneys, and liver were harvested 24 hours after injection for fibrotic kidney experiments and 48 hours after injection for fibrotic lung experiments. Fluorescence intensity was measured via IVIS (images in Supplemental Figure 5) and the normalized fluorescence units as % of total fluorescence was calculated for healthy and fibrotic kidney and lungs. N ranges from 2 to 4. * = statistical significance of P < 0.05 between fluorescence of fibrotic organs vs fluorescence of healthy organs, Student’s t-test.

In this fibrosis targeting study, we observed unexpectedly that α-FN-EDA Fab targeted to fibrotic kidneys, but not to fibrotic lungs (Figure 1). This result may be due to differences in the amount fibronectin-EDA present in the kidney and lung fibrosis models used in the study. Further investigation outside of the scope of this study is needed to explore this hypothesis. While α-FN-EDA appears not to be suitable for targeting bioactive macromolecules to fibrotic lungs, it may be used in the future as a drug carrier in kidney fibrosis. However, in the same targeting study, VWF-A3 displayed greater accumulation in both fibrotic kidneys and lungs compared to the healthy organs (Figure 1). This likely occurs due to high levels of exposed collagen I and III present in both disease models [9,10]. We also evaluated whether CBP-micelles could target exposed collagen in fibrotic kidneys and lungs (Figure 1). We observed that CBP-micelles preferentially accumulated in fibrotic kidneys with total fluorescence units exceeding 20% in both mouse models (Figure 1). Scrambled CBP-micelles also accumulated in the fibrotic tissues, but to a lesser extent than the actively targeted CBP-micelles (Figure 1). The observed targeting of scrambled CBP-micelles, especially in the lung fibrosis model, is likely due to the physical effect of the passive targeting of nanoparticles to tissues with increased vascular permeability [35, 36]. Because VWF-A3 and CBP-micelles could successfully target to both fibrotic kidneys and lungs, we selected these delivery systems as carriers of IL-10 and rapamycin, respectively.

### 2.3. Collagen-Targeting IL-10 Treatment Suppresses Kidney and Lung Fibrosis

We first recombinantly fused the VWF-A3 to IL-10 (A3-IL-10) and produced this protein in human embryonic kidney cells (Supplementary Figure 6) [26, 27, 37, 38]. To prolong the half-life of the collagen-targeted IL-10, we incorporated mouse serum albumin (A3-SA-IL-10) (Supplementary Figure 6) as described previously [37]. Following affinity and size exclusion chromatography, the purity of A3-IL-10 and A3-SA-IL-10 variants was confirmed by SDS-PAGE (Supplementary Figure 9A). Affinity measurements were then performed to confirm the ability of IL-10 variants to bind to IL-10 receptor alpha (Supplementary Figure 9B) and collagen I and III (Supplementary Figure 10A, B).

To determine if collagen targeting A3-IL-10 and A3-SA-IL-10 could treat kidney fibrosis more effectively than unmodified IL-10, we administered these therapies in a mouse model of kidney fibrosis. In this model, fibrosis was induced in the left kidney via UUO ligation and 7 days post UUO, mice were intravenously injected with 20 μg (or molar equivalent) of IL-10, A3-IL-10, and A3-SA-IL-10. Mice were then sacrificed 14 days post UUO, and both injured and fibrotic kidneys were resected, fixed, sectioned, and stained for collagen I using immunohistochemistry. The extent of fibrosis in uninjured (Figure 2A), untreated fibrotic (Figure 2B), and IL-10 treated fibrotic kidneys (Figure 2C-E) was then assessed by quantifying the ratio of fibrotic (collagen I high) to healthy (collagen I low) tissue area (Figure 2F). While unmodified IL-10 treatment slightly reduced extent of kidney fibrosis, only collagen-targeted A3-IL-10 and A3-SA-IL-10 significantly reduced the disease area ratio. Furthermore, the reduction of fibrotic to healthy tissue area ratio in mice treated with A3-SA-IL-10 was significantly lower than mice treated with unmodified IL-10.

**Figure 2.**
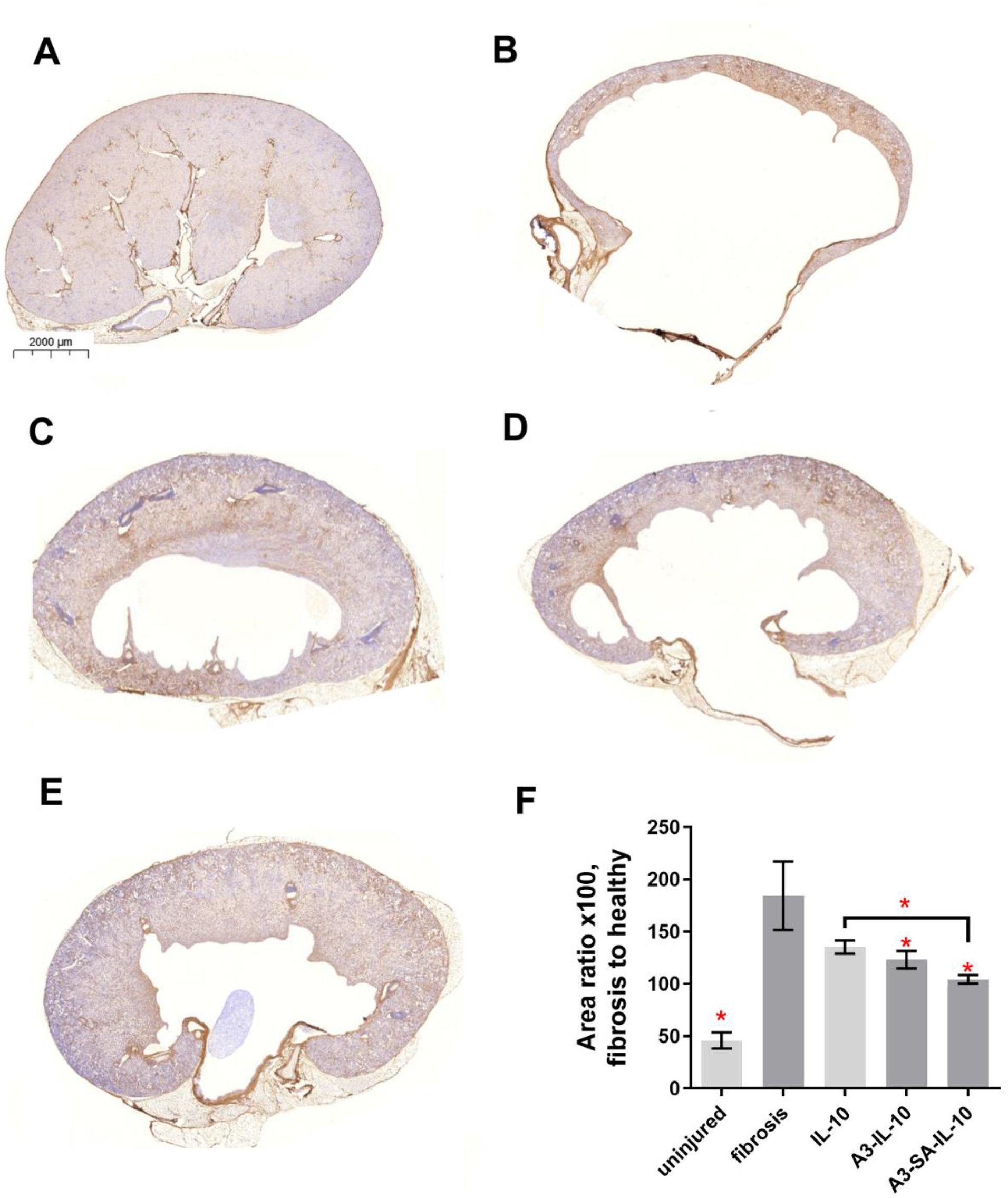
A single injection of 20 µg of IL-10-A3 or A3-SA-IL-10 rescues the fibrotic damage from UUO insult to mouse kidneys. The descending ureter of the left kidney was surgically ligated, and 20 µg (or molar equivalent) IL-10, IL-10-A3, or IL-10-A3-MSA was injected i.v. after 7 days. Kidneys were resected 14 days post UUO insult (7 days post antibody injection), mounted, and assessed via immunohistochemistry for collagen I. The amount of positive IHC staining was compared to the overall amount of kidney tissue, per image. Representative images for (A) uninjured kidneys, (B) fibrotic kidneys, and fibrotic kidneys treated with (C) IL-10, (D) A3-IL-10, and (E) A3-SA-IL-10.(F) Area ratio (x100) of fibrotic to healthy tissue calculated based on histology images. N=6. * = statistical significance of P < 0.05, ANOVA vs fibrosis control, Welch’s correction. Comparison between IL-10 and A3-SA-IL-10 is Student’s t-test.

We next evaluated the efficacy of collagen targeted A3-IL-10 and A3-SA-IL-10 in a mouse model of lung fibrosis. For this purpose, we insulted the lungs with bleomycin and treated the mice by intravenously injecting 10 μg (or molar equivalent) of IL-10, A3-IL-10 and A3-SA-IL-10 at 7, 10, 14 and 17 days post-bleomycin insult (Figure 3A-H). Because bleomycin injury is associated with weight loss in mice [19], we weighed the mice, but noted no significant changes in weight between the experimental groups (Figure 3I). The mice were then euthanized 21 days post insult. A hydroxyproline assay was performed on the right lung lobe to quantify the amount of collagen, which is positively correlated with ongoing fibrotic disease (Figure 3J). From this assay, we determined that A3-IL-10 significantly reduced the amount of collagen in the right lobe of the fibrotic lungs, whereas unmodified IL-10 had no significant impact on this metric (Figure 3J). The left lung lobes were also resected, fixed, sectioned, and stained for Masson’s trichrome, a dye used to visualize collagen present in fibrotic tissue sections. The stained histology images were blindly scored using the Ashcroft scale, an established method for quantifying the extent of lung fibrosis (Figure 3A-H). This assessment confirmed the results of the hydroxyproline assay. A3-IL-10 treatment significantly reduced Ashcroft score compared fibrotic untreated and unmodified IL-10 treated mice (Figure 3A-H). Taken together, these results suggest that collagen targeting via recombinant fusion of VWF-A3 to IL-10 improves the therapeutic efficacy of IL-10 in mouse models of kidney and lung fibrosis. In the kidney fibrosis model, the addition of half-life prolonging serum albumin to the A3-IL-10 construct further improved efficacy, whereas in the lung fibrosis model, this modification did not show any additional benefit over A3-IL-10. The differences in efficacy due to serum albumin fusion may have been influenced by the different dosing regimens in the two mouse models. In the lung fibrosis model, the mice were dosed more frequently with the IL-10 variants, which may not have taken advantage of the prolonged half-life of the serum albumin fused molecule. Future studies can be performed to further optimize the dose of collagen-targeted therapies.

**Figure 3.**
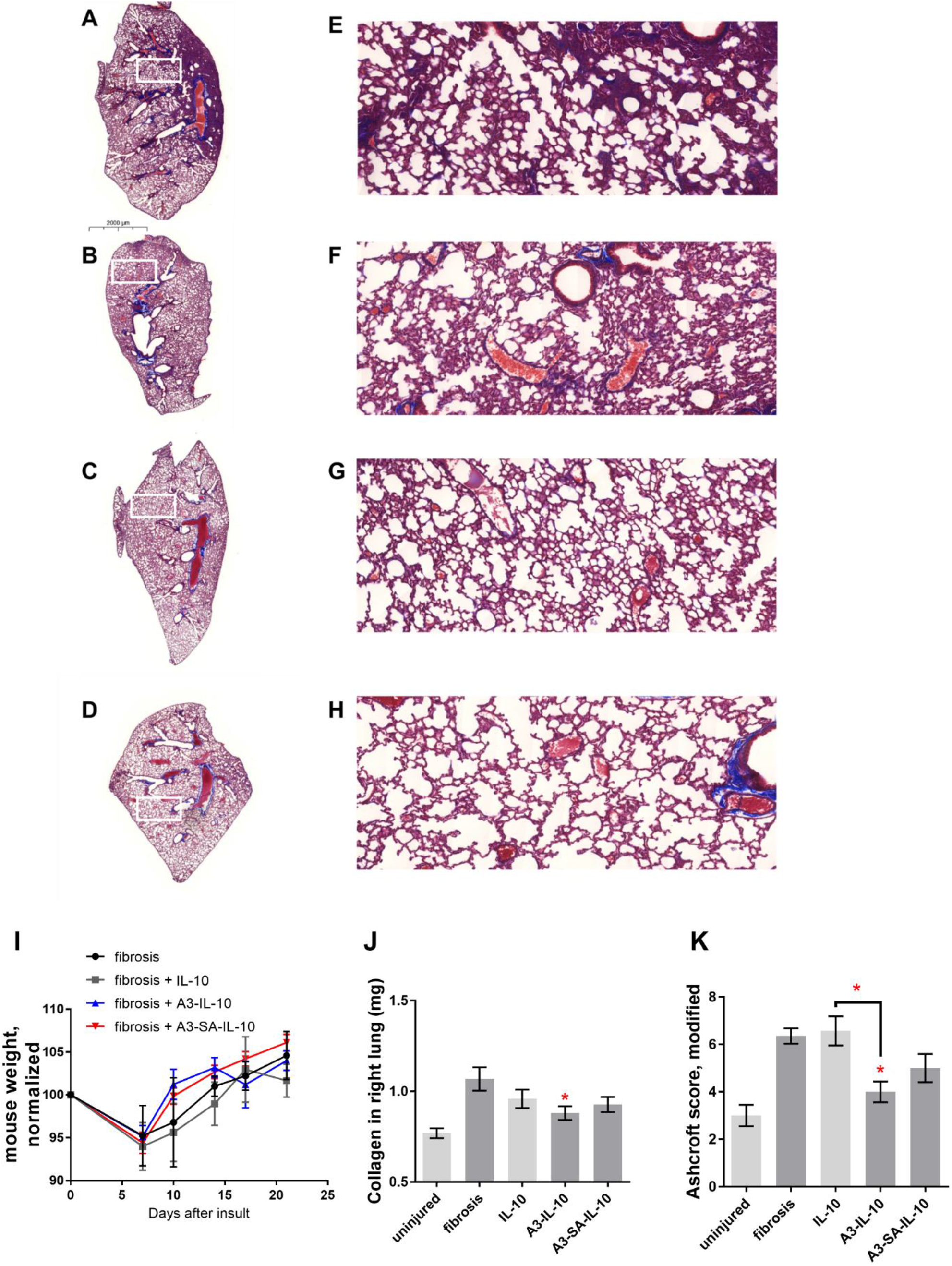
A3-IL-10 rescues the fibrotic damage in mouse model of lung fibrosis. Lungs insulted with 75 μg bleomycin; 1 week following instillation mice were left untreated (A,E) or injected i.v. with a total of 40 µg (or molar equivalent) in 4 injections of 10 µg on days 7, 10, 14 and 17 post bleomycin insult with either (B, F) IL-10, (C, G) A3-IL-10, (D,H) A3-SA-IL-10. Insets are pictured and the size bar is 2000 µm. (I) Mouse body weights after bleomycin insult and treatment. (J) Collagen content from the right, multi-lobed lung assessed by hydroxyproline assay. (K) Blinded Ashcroft scoring. N = 8. * = statistical significance of P < 0.05, significance vs fibrotic lungs calculated by 2-way ANOVA (with Fisher’s LSD post-test). Comparison between IL-10 and A3-IL-10 is Student’s t-test.

### 2.4. Rapamycin Encapsulated Collagen-Targeting Micelles Suppress Kidney and Lung Fibrosis

Having demonstrated that the recombinant fusion of collagen targeting domains to the biologic drug IL-10 improves its therapeutic efficacy in mouse models of fibrosis, we next sought to evaluate whether the encapsulating the small molecule drug rapamycin in collagen binding peptide-conjugated micelles could also improve efficacy in these models. For this purpose, we induced kidney fibrosis in the left kidney via UUO ligation. Then, we injected 3 mg/kg of rapamycin as a free molecule or encapsulated in CBP-micelles (rapamycin-CBP-micelles) at 7-, 9-, 11-and 13-days post disease induction (Figure 4). On day 14, mice were sacrificed, and tissues were processed and analyzed as described above. The extent of fibrosis in uninjured (Figure 4A), untreated fibrotic (Figure 4B), free rapamycin treated fibrotic (Figure 4C) and rapamycin-CBP-micelles treated fibrotic (Figure 4D) kidneys was then assessed by quantifying the ratio of fibrotic (collagen I high) to healthy (collagen I low) tissue area (Figure 4E). While both rapamycin treatments significantly reduced kidney fibrosis, rapamycin-CBP-micelles treatment showed the greatest reduction in collagen I fibrotic-to-healthy disease area ratio. The induction of fibrosis has been reported to alter blood chemistry parameters [39], we compared the levels of markers of liver damage and kidney damage (alanine aminotransferase (ALT), albumin, amylase, aspartate aminotransferase (AST), total bilirubin, blood urea nitrogen (BUN), calcium, creatinine, creatine kinase, total bilirubin, and uric acid), in the serum of uninjured, fibrotic and rapamycin-treated fibrotic mice. Of these, rapamycin treatment changed the levels of alanine aminotransferase (ALT) (Supplementary Figure 11A), aspartate aminotransferase (AST) (Supplementary Figure 11B) and total bilirubin (Supplementary Figure 11C), In this study, we observed that kidney fibrosis induction reduces ALT, AST and total bilirubin levels in the serum (Supplementary Figures 11A-C), which is consistent with published reports on the response of ALT, AST, and bilirubin to fibrosis [40]. While treatment of fibrotic mice with free rapamycin did not affect biochemistry markers, we observed that treatment with collagen-targeting rapamycin CBP-micelles restored ALT, AST and total bilirubin levels to those of uninjured mice (Supplementary Figures 11A-C).

**Figure 4.**
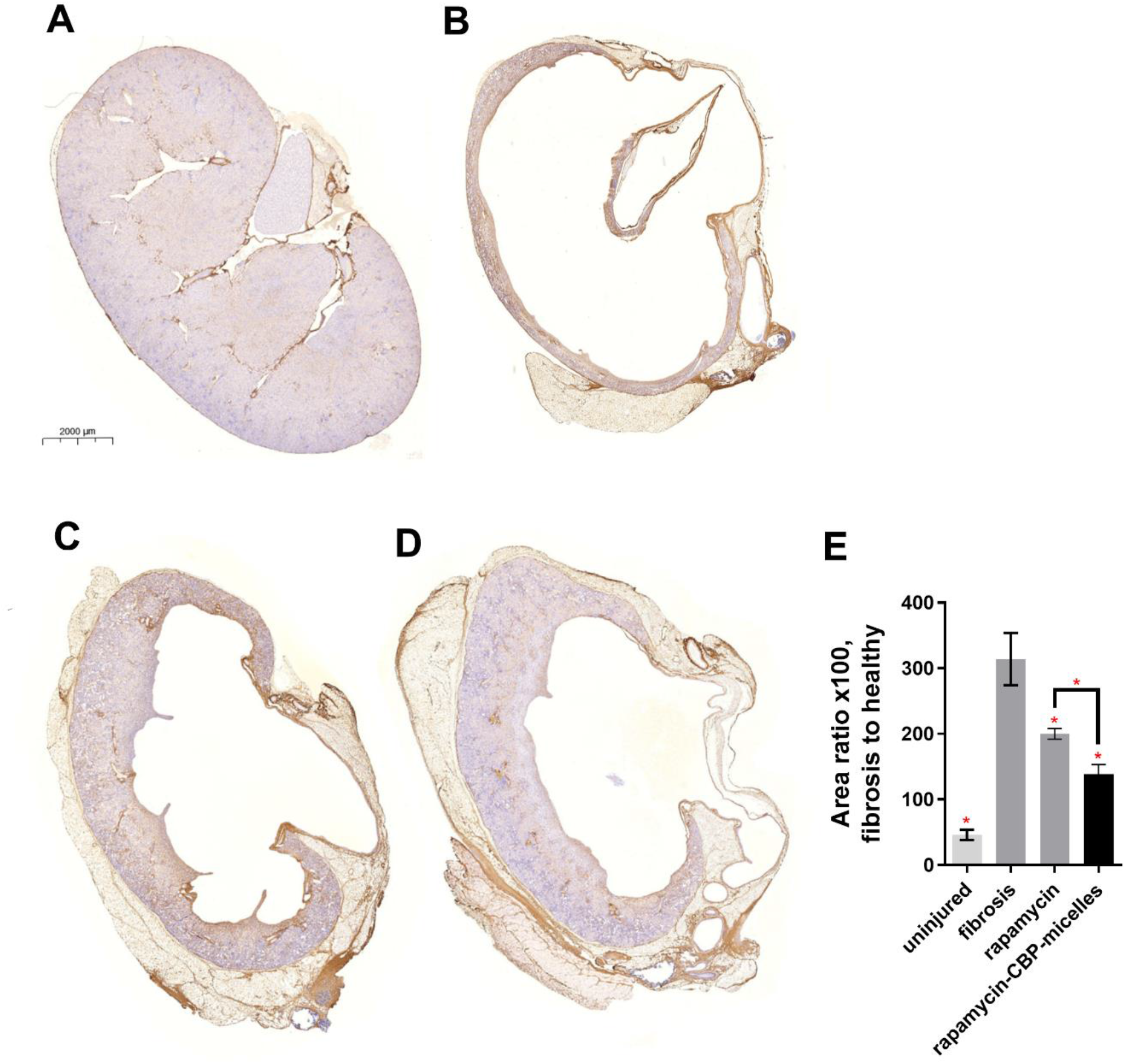
Rapamycin and rapamycin-CBP-micelle treatments rescue the fibrotic damage from UUO insult to mouse kidneys. The descending ureter of the left kidney was surgically ligated, and 3 mg/kg (or molar equivalent) rapamycin and rapamycin-CBP-micelles was injected i.v. at 7, 9, 11, and 13 days post UUO ligation. Kidneys were resected 14 post UUO insult, mounted, and assessed via immunohistochemistry for collagen I. Representative images for (A) uninjured kidneys, (B) fibrotic kidneys, (C) and fibrotic kidneys treated with (C) rapamycin, and (D) rapamycin-CBP-micelles. (E) The amount of positive IHC staining was compared to the overall amount of kidney tissue, per image. N=6. * = statistical significance of P < 0.05, ANOVA vs fibrosis control, Welch’s correction. Comparison between rapamycin and rapamycin-CBP-micelles is Student’s t-test.

**Figure 5.**
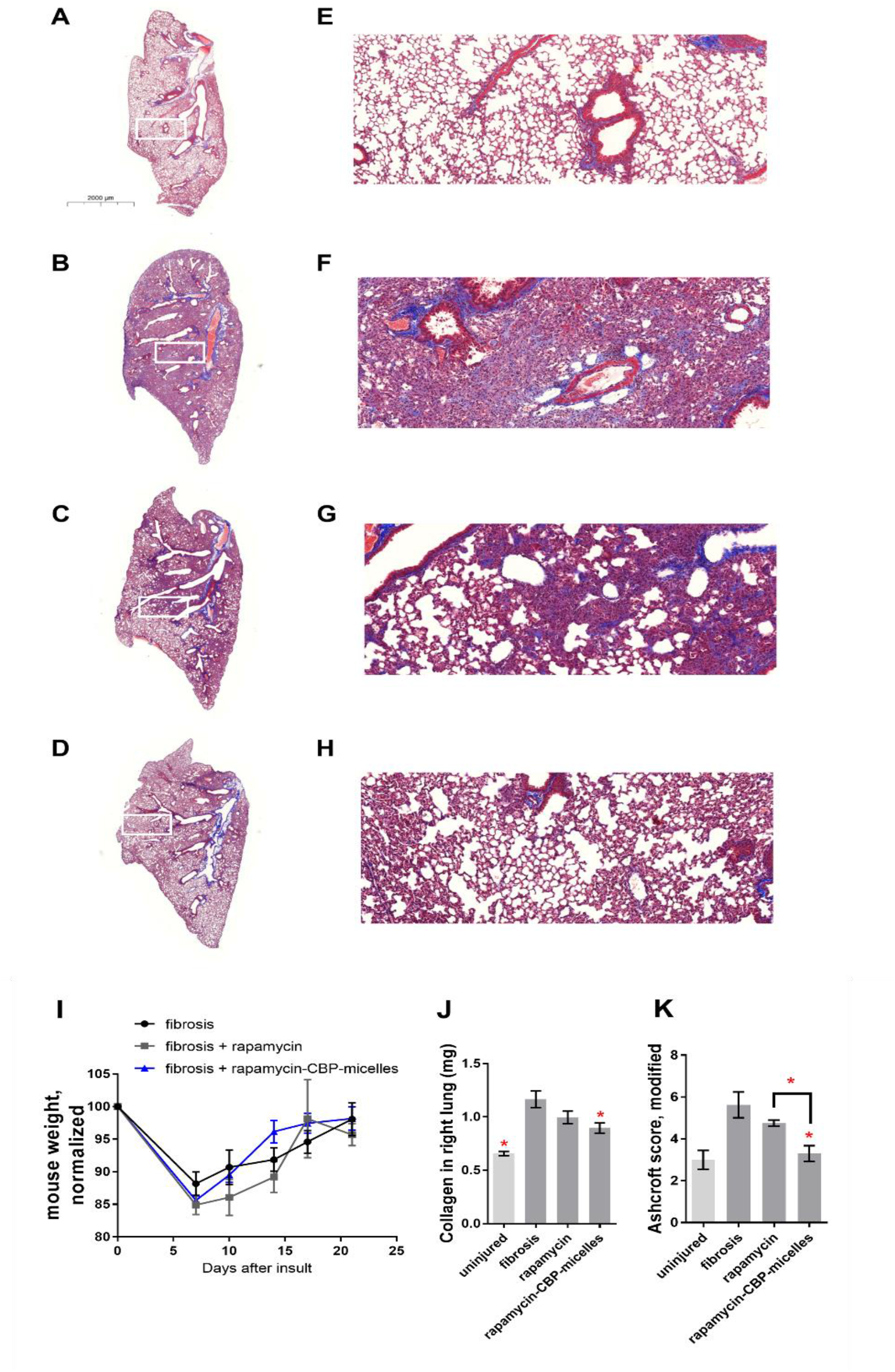
Rapamycin and rapamycin-CBP-micelle treatments rescue the fibrotic damage in a mouse model of lung fibrosis. Lungs were insulted with 75 μg bleomycin and 1 week later mice were left untreated (B,F) or injected i.v. with 3 mg/kg of free rapamycin (C,G) or rapamycin-CBP-micelles (D,H) on days 7, 9, 11, 13, 15, 17 and 19. Healthy lungs (A,E) were imaged for reference. Insets are pictured and the size bar is 2000 microns. (I) Mouse weights after bleomycin insult and treatment. (J) Collagen content from the right, multi-lobed lung assessed by hydroxyproline assay. (K) Blinded Ashcroft scoring. N = 7 for untreated fibrosis, N = 4 for rapamycin-treated fibrosis, and N= 6 for rapamycin-CBP-micelles treated fibrosis. * = statistical significance of P < 0.05, significance vs fibrotic lungs calculated by 2-way ANOVA (with Fisher’s LSD post-test). Comparison between rapamycin and rapamycin-CBP micelles is Student’s t-test.

After demonstrating the efficacy of rapamycin formulated in collagen targeting CBP-micelles in kidney fibrosis, we next sought to evaluate its efficacy in a mouse model of lung fibrosis. For this purpose, we induced lung fibrosis via bleomycin instillation into the lungs. Fibrotic mice were then treated with intravenous administration of 3 mg/kg of rapamycin as a free molecule or encapsulated in CBP-micelles (rapamycin-CBP-micelles) at 7, 9, 11, 13, 15, 17 and 19 days post disease induction (Figure 6A-H). Throughout the experiment, we observed no significant changes in weight between the experimental groups (Figure 6I). The mice were euthanized 21 days post insult and the lungs were processed and analyzed as described above. Based on the hydroxyproline assay, the administration of rapamycin CBP-micelles significantly reduced the amount of collagen in the right lobe of the fibrotic lungs, whereas free rapamycin treatment had no significant impact on this metric (Figure 6J). Similarly, blinded analysis of Masson’s trichrome staining of lung histology sections (Figure 6A-H) showed that rapamycin CBP-micelles treatment reduced Ashcroft score of fibrotic left lungs to healthy lung levels and that the score was significantly lower compared to free rapamycin treated fibrotic mice (Figure 6K). In this study, we also analyzed blood chemistry markers of uninjured, untreated fibrotic and rapamycin-treated fibrotic mice. For this purpose, we quantified the concentration of markers of lung, heart and liver damage (alanine aminotransferase (ALT), albumin, amylase, aspartate aminotransferase (AST), total bilirubin, blood urea nitrogen (BUN), calcium, creatinine, creatine kinase, CO_2_, total bilirubin, and uric acid) in the serum,. While rapamycin-CBP micelles did not significantly improve CO_2_ levels in the bloodstream (Supplementary Figure 12A), treatment with rapamycin-CBP-micelles restored creatine kinase (Supplementary Figure 12B), ALT (Supplementary Figure 12C) and AST (Supplementary Figure 12D) levels to those of uninjured mice. Taken together, these results suggest that collagen targeting of rapamycin via its encapsulation in CBP-micelles improves the therapeutic efficacy of rapamycin in mouse models of kidney and lung fibrosis. Therapeutic outcomes could be further improved by optimizing the dose of rapamycin administered and the treatment regimen. The dose of rapamycin used in this study (3 mg/kg) is lower than doses previously used in the treatment of kidney [24] and lung [25] fibrosis. The dose used, administered every other day, is also comparable to the lowest average daily dose found in the relevant literature (1.5 mg/kg) [41]. Our CBP-micelle technology can also be adapted to formulate other anti-fibrotic water-insoluble molecules currently in development to improve their solubility, pharmacokinetics, toxicity and efficacy.

## 3. Conclusion

Fibrosing diseases are refractory to treatment, and currently available therapies only mildly reduce the symptoms of disease. A major goal of the ongoing fibrosis research is to identify new clinical treatments that are able to reverse damage and restore function of affected organs. In this work, we have demonstrated that novel collagen-binding protein-based and nanoparticle-based drug delivery systems can be harnessed to target systemically administered bioactive molecules to the leaky and collagen-rich fibrotic organs. We have engineered and validated formulations of two candidate drugs: IL-10, an anti-inflammatory biologic, and rapamycin, an anti-fibrotic small molecule water-insoluble drug. In concert, our studies demonstrated improved efficacy of A3-IL-10/A3-SA-IL-10 and rapamycin-CBP-micelles in mouse models of kidney and lung fibrosis. These therapies are particularly promising because each of IL10 (Supplementary Figure 13), IL10 receptor (IL10R, Supplementary Figure 14), and mTOR (Supplementary Figure 15) are downregulated in idiopathic pulmonary fibrosis (IPF, data from IPF cell atlas [17]).

Our novel therapies outperformed the unmodified drugs in reversing the phenotypic changes in fibrosis-affected organs and outperformed the unmodified drugs. The technologies presented in this work can be easily adapted in the future to deliver other biologics as well as water-insoluble medicines. Taken together, our findings indicate that collagen-targeted approaches can be used to develop versatile the next generation anti-fibrotic therapies with high clinical potential.

## 4. Experimental Section

### Study design

This study was designed to screen for fibrosis-targeting agents from several collagen-targeting proteins, peptide-decorated micelles and antibodies. Group size was selected taking into account the reported variability in mouse kidney and pulmonary fibrosis models. To eliminate cage effects, mice were randomized into treatment groups within a cage. To ensure reproducibility, treatment was performed by multiple researchers over the course of this study. Kidneys and lungs were also resected by multiple researchers, and blinded scoring was performed on the lung fibrosis histology images.

### Production of α-Fibronectin-EDA Fab

The sequence encoding the anti-fibronectin-EDA Fab (α-FN-EDA Fab) was synthesized and subcloned into the mammalian expression vector pSecTag A. Suspension-adapted HEK-293F cells were routinely maintained in serum-free FreeStyle 293 Expression Medium (Gibco). On the day of transfection, cells were inoculated into fresh medium at a density of 1 × 10^6^ cells/mL, and then 2 µg/mL plasmid DNA, 2 µg/ml linear 25 kDa polyethylenimine (Polysciences), and OptiPRO SFM medium (4% final concentration, Thermo Fisher) were sequentially added. The culture flask was agitated by orbital shaking at 135 rpm at 37°C in the presence of 5% CO_2_. 7 days after transfection, the cell culture medium was collected by centrifugation and filtered through a 0.22 µm filter. Next, the cell culture medium was loaded onto a HiTrap MabSelect 5 mL column (GE Healthcare), using an ÄKTA Pure 25 (GE Healthcare). After washing the column with PBS, protein was eluted with 0.1 M sodium citrate (pH 3.0). Purification was carried out at 4°C. The α-FN-EDA Fab was verified as >90% pure by SDS-PAGE (performed on 4-20% gradient gels (Bio-Rad)).

### Production and purification of recombinant A3-IL-10 and A3-SA-IL-10

Recombinant VWF-A3 cytokine fusion proteins were produced as previously described [18-20]. Briefly, the sequences encoding the A3 domain of VWF (A3), a (GGGS)_2_ flexible linker, mouse serum albumin (SA), a (GGGS)_2_ flexible linker and mouse IL-10 (A3-SA-IL-10) or A3, (GGGS)_2_ and mouse IL-10 (A3-IL-10) were synthetized and subcloned into the mammalian expression vector pcDNA3.1(+) by GenScript. To enable affinity-based purification, a sequence encoding (His)_6_ was added to the carboxy terminus of each protein. The amino acid sequences of the proteins are shown in Supplementary Figure 6. Suspension-adapted HEK293F cells were maintained in serum-free Free Style 293 Expression Medium (Gibco). On the day of transfection, HEK cells were resuspended in fresh medium at a density of 10^6^ cells mL^-1^. A concentration of 2 μg mL^-1^ plasmid DNA and 2 μg mL^-1^ plasmid DNA linear 25kDa polyethyleneimine (Polysciences) dissolved in OptiPRO serum free medium (Thermo Fisher) at 4% final volume were added sequentially to the suspension cells. After 7 days of culture, cell supernatants were collected, centrifuged at 4,000 rpm for 10 minutes at 4°C, and filtered through a 0.22 μm filter. Proteins were purified via affinity chromatography and size exclusion chromatography using an ÄKTA pure 25 (Cytiva) as described previously [37]. Protein purity was assessed by SDS-PAGE (Supplementary Figure 9A) [18-20]. The purified proteins were then tested for endotoxin using the HEK-Blue TLR4 reporter cell line, and the endotoxin levels were confirmed to be below 0.01 EU mL^-1^ [18-20]. Protein concentration was determined via absorbance at 280 nm using a NanoDrop spectrophotometer (Thermo Scientific).

### Detection of A3-IL-10 and A3-SA-IL-10 binding to collagens

Surface plasmon resonance (SPR) measurements were performed using the Biacore X100 SPR system (Cytiva) as described previously [18]. To measure the affinity of A3-IL-10 and A3-SA-IL-10 for collagen proteins, recombinant human type I or type III collagen (Millipore Sigma) were immobilized on a CM5 sensor chip (Cytiva) by standard amine coupling method for around 1500 resonance units (RUs) and blocked with ethanolamine according to the manufacturer’s instructions. A reference cell was also blocked with ethanolamine. Binding assays were carried out at room temperature and the *K*_*d*_ values of A3-IL-10 and A3-SA-IL-10 were determined by fitting the 1:1 Langmuir binding model to the data using BIAevaluation software (Cytiva) and are displayed in Supplementary Figure 10A, B.

### Detection of A3-IL-10 and A3-SA-IL-10 binding to IL-10 receptor

An ELISA was performed to determine the affinity of IL-10 variants for IL-10 receptor as described previously for A3-cytokine variants [20]. 96-well ELISA plates (Greiner Bio One) were coated with 4 μg ml^-1^ recombinant mouse IL-10 receptor alpha (R&D Systems) overnight at 4°C. The wells were then washed in PBS with 0.05% Tween 20 (PBS-T) and blocked with 2% BSA in PBS-T for 1 hour at room temperature. 0 to 4 μg ml^-1^ A3-IL-10 or A3-SA-IL-10 (wild type IL-10 molar equivalent) were added to the wells and incubated for 2 hours at room temperature. After three washes with PBS-T, wells were then incubated for 1 hour at room temperature with biotinylated antibody against IL-10 (Invitrogen) and then incubated with HRP-conjugated streptavidin (Invitrogen) for 30 minutes at room temperature. After five washes, the bound A3-IL-10 and A3-SA-IL-10 were detected by tetramethylbenzidine substrate measurement of absorbance at 450 nm with subtraction of 570 nm as described previously [20]. The apparent K_D_ values were obtained by nonlinear regression analysis in Prism software (v9, GraphPad Software assuming one-site specific binding) and are displayed in Supplementary Figure 9B.

### Synthesis of pEG_114_-pPhe_20_ copolymers

Both methoxy-pEG_114_-pPhe_20_ and N_3_-pEG_114_-pPhe_20_ polymers were prepared by ring-opening polymerization of L,D-Phenylalanine N-carboxyanhydride (Phe-NCA; made in house from L,D-Phenylalanine and triphosgene) using amino-terminated methoxy-pEG_114_-NH_2_ and N_3_-pEG_114_-NH_2_, respectively, as initiators [21]. Briefly, Phe-NCA (50 mg, 0.25 mmol) and respective pEG_114_-NH_2_ (50 mg, 0.01 mmol) were added to anhydrous dimethylformamide: tetrahydrofuran solution (DMF:THF 1:1 v/v, 2 mL) under a nitrogen atmosphere for further reaction time of 24 h at 35°C (Supplementary Figure 1A). The copolymers were purified by multiple precipitation in diethyl ether and hexanes, and dried under vacuum for further use. The yield of polymerization was 66% for methoxy-pEG_114_-pPhe_20_ and 71% for N_3_-pEG_114_-pPhe_20_. The polymers were characterized using ^1^H NMR and MALDI-TOF to confirm the composition and purity (Supplementary Figure 1B, C).

#### Determination of critical micelle concentration (CMC)

The CMC of block copolymer pEG_114_-pPhe_20_ was determined by fluorescence and using pyrene as a probe based on previously reported procedures [146]. Briefly, micellar dispersions of pEG_114_-pPhe_20_ were prepared at various concentrations (0.1 -0.13 mM) with a fixed concentration of pyrene of 0.6 µM. The excitation spectra of pyrene from 300 to 360 nm were monitored as 390 nm for each dilution using Horiba Scientific Fluoromax 4 fluorimeter. CMC was determined by plotting the ratio of the intensities at 335 and 332 nm (I_3_/I_1_) versos concentration of pEG_114_-pPhe_20_.

#### *In vitro* release kinetic study

To study the kinetics of drug release from pEG_114_-pPhe_20_ micelles, model drug/dye Rhodamine B was first encapsulated in pEG_114_-pPhe_20_. The free dye was removed using Zeba desalting columns with MWCO of 7kDa. The release kinetic was assessed in vitro by employing the dialysis method [152]. Equal amount of Rhodamine B, either as a free dye or formulated within pEG_114_-pPhe_20_ micelles, was placed in a tube capped with a dialysis membrane of MWCO of 10kDa. The formulations were dialyzed against 10x volume of PBS buffer. The profiles were obtained by periodically withdrawing aliquots of the dialys buffer to measure the amount of released Rhodamine B via fluorescence spectroscopy (ex/em 550/600) using Horiba Scientific Fluoromax 4 fluorimeter.

### Synthesis of AF647-labeled pEG_114_-pPhe_20_ copolymers

Alexa Fluor™ 647 Succinimidyl ester (AF647-NHS, Invitrogen) was used to label pEG_114_-pPhe_20_ copolymers for fluorescence imaging biodistribution study evaluation. Briefly, AF647-NHS (1 mg) was added to pEG_114_-pPhe_20_ (25 mg) in anhydrous dichloromethane (1 mL) and stirred for 24 hours. The pEG_114_-pPhe_20_-AF647 conjugates were purified by precipitation in diethyl ether.

### Synthesis of peptide-linker azide-reactive conjugates

Both collagen binding peptide (CBP, sequence: CNNNLHLERL, Genscript) and scrambled sequence peptide (SCR, sequence: CLLNEHLNNR, Genscript) were modified with heterobifunctional Sulfo-DBCO Maleimide linker (Click Chemistry Tools) to arm the peptide with an azide-reactive group (dibenzocyclooctyne group, DBCO). Briefly, Sulfo-DBCO Maleimide linker (0.44mg, 1.0 eq) was added to peptide (1 mg, 1.1 eq) solution in DMSO and allowed to react overnight (Supplementary Figure 3). The peptide-linker was used crude without further modification.

### Formulation and characterization of peptide-micelle conjugate nanoparticles

Copolymer micelles were formed by rehydration of the polymer film and subsequent sonication of resulted suspension (Supplementary Figure 3). Briefly, methoxy-pEG_114_-pPhe_20_-AF647 (9.5 mg) and N_3_-pEG_114_-pPhe_20_-AF647 (0.5 mg) solution (to ensure 5%mass of azide-terminated polymer) in tetrahydrofuran was transferred to a 1.5 mL glass vial and the polymer film was formed under nitrogen gas flow. Next, phosphate buffered saline (PBS, 1 mL) and the solution was sonicated for 45 minutes until almost-clear uniform suspension was formed. The suspension was then filtered with 0.22 µm PVDF syringe filter (ThermoFisher) and peptide-linker solution (at 1.1:1 linker:N_3_ ratio) was added. The mixture was allowed to react at room temperature for 6 hours (Supplementary Figure 3) and then was dialyzed against PBS using Slyde-A-Lyzer™ Dialysis Cassette (7K MWCO, 3mL, ThermoFisher) to remove unreacted peptide-linker and any remaining free fluorescent dye (AF647). Obtained micelle-peptide nanoparticle solutions: 5%N_3_-pEG_114_-pPhe_20_, CBP-pEG_114_-pPhe_20_, and SCR-pEG_114_-pPhe_20_ were characterized using Dynamic Light Scattering (DLS) (Supplementary Figure 4A) and cryogenic electron microscopy (cryo-EM) (Supplementary Figure 4B) to determine their size, zeta potential and morphology.

### Formulation and characterization of rapamycin-loaded CBP-micelles

Rapamycin (MedChemExpress) (5 mg) in tetrahydrofuran (200 µL) was added in to methoxy-pEG_114_-pPhe_20_-AF647 (9.5 mg) and N_3_-pEG_114_-pPhe_20_-AF647 (0.5 mg) in tetrahydrofuran (300uL) in 1.5 mL glass vial. The polymer-drug film was formed under nitrogen gas flow. Next, phosphate buffered saline (PBS, 1 mL) and the solution was sonicated for 45 minutes until almost-clear uniform suspension was formed. The suspension was then filtered with 0.22 µm PVDF syringe filter (ThermoFisher) and peptide-linker solution (at 1.1:1 linker:N_3_ ratio) was added. After reacting for 6 hours, the formulation was ready for use. The concentration of rapamycin was determined by UV-VIS at 290 nm, using a standard curve method.

### Formulation of rapamycin for intravenous injection

Free rapamycin saline solution was prepared by diluting a stock solution of rapamycin in DMSO:PEG300:Tween80:saline (5:40:5:50 v/v) to a desired concentration.

### Production of fluorescently labeled antibodies

α-FN-EDA Fab, and VWF-A3 were fluorescently labeled with Cy7 using sulfo-Cy7 *N*-hydroxysuccinimide ester (Lumiprobe) according to the manufacturer’s instruction, as previously described [22]. Unreacted Cy7 was removed by dialysis against PBS.

### In vivo biodistribution study

An *in vivo* biodistribution study was conducted as previously described [22], with minor adjustments. 7 days following bleomycin insult or UUO surgery, 50 µg (or molar equivalent) of Cy7-α-FN-EDA Fab, Cy7-VWF-A3, and Cy7 were injected via tail vein. 24 hours later or 48 hours later (Figure 1), heart, lungs, spleen, kidneys, and liver were resected and imaged via IVIS (Xenogen) under the following conditions: *f*/stop, 2; optical filter excitation, 710 nm; emission, 780 nm; exposure time, 5 seconds; small binning.

### Bleomycin induced pulmonary fibrosis model

Male and female mice were acquired at 8 weeks of age (Jackson laboratories, Bar Harbor, ME). Mouse lungs were instilled with 0.075 units bleomycin (75 µg, Fresenius Kabi, Switzerland) suspended in endotoxin-free PBS, as previously described [22]. Briefly, mice were anesthetized via isoflurane inhalation (2%). Mice were then placed upright on an angled surface and a 200 µL narrow pipet was placed at the entrance of their throat. 50 µL of bleomycin/PBS was dispensed to the entrance of the throat, and mice were allowed to inhale. Mice were then weighed and placed on a heating pad to recuperate.

Following bleomycin insult, mice were injected via tail-vein with 10 µg (or molar equivalent) of IL-10, A3-IL10, or A3-SA-IL-10 on 7, 10, 14, and 17 days after bleomycin insult. Another group of mice were injected with 3 mg/kg of rapamycin (or molar equivalent) or rapamycin-CBP-micelles via tail-vein injection 7, 9, 11, 13, 15, 17 and 19 days following bleomycin insult.

Mice were euthanized at 21 days post insult via injection of euthasol (Covetrus, Portland, ME) instead of CO_2_ inhalation to prevent the potential damage of the lungs.

### Lung resection and fibrosis scoring

Lungs were harvested, and the right and left lobes were separated. The left lobe was fixed in 4% paraformaldehyde overnight, mounted in paraffin, sectioned into 5 mm slices, and stained using Masson’s trichrome. Stained lungs were scanned at high resolution using a CRi Panoramic SCAN 40x Whole Slide Scanner (Perkin-Elmer, Waltham, MA), and were read for fibrosis using a modified Ashcroft method, as previously described [22]. Lungs were read unlabeled by another researcher uninvolved with animal treatment.

The right lobe of the lung was frozen, and dehydrated using a tissue lyophilizer (Labconco, Kansas City, MO), weighed, and was assessed for collagen content by hydroxyproline assay [23]. Briefly, dried lungs were digested in 6N HCl/PBS at 100°C for 24 hours. Supernatant from this digestion was added to 96 well plates and treated sequentially with chloramine-T solution and Ehrlich’s solution at 65°C for 15 minutes to facilitate the color change reaction. Color was read at 561 nm. Quantification was provided by use of a hydroxyproline (Sigma) dilution series, which was transformed into a standard curve.

### Kidney unilateral ureteral obstruction (UUO) fibrosis model

UUO surgery was performed as previously described [22], with adjustments. Briefly, mice were anesthetized via 2% isoflurane inhalation, and injected with meloxicam (1 mg/kg), buprenorphine (0.1 mg/kg) in a saline solution, subcutaneously. Mice were laid on their right side and an abdominal incision used to visualize the left ureter. The left ureter was ligated in the middle section of the ureter with two ties (2 mm apart) using 7-0 silk sutures. Peritoneum was then closed with 5-0 vicryl sutures and skin closed with 5-0 nylon sutures.

One week following UUO ligation, mice were injected via tail-vein with 10 µg (or molar equivalent) of IL-10, A3-IL-10, or A3-SA-IL-10. 7, 10, 14, and 17 days after UUO insult. Another group of mice were injected with 3 mg/kg (or molar equivalent) of rapamycin or rapamycin-CBP-micelles via tail-vein injection 7, 9, 11, 13, 15, 17 and 19 days following UUO insult. Two weeks post UUO, and one week post injection, mice were sacrificed via CO_2_ inhalation, and their kidneys harvested. At this point, the placement of UUO ligation was confirmed for each mouse.

### Assessment of fibrosis in kidneys

Kidney fibrosis was assessed as previously described [22]. Briefly, right (healthy) and left (fibrotic) kidneys were placed in 4% PFA for 24 hours, mounted in paraffin, sectioned into 5 mm full kidney slices, and stained using immunohistochemistry (IHC) for collagen I (1:4000, polyclonal rabbit, lifespan biosciences, Seattle WA) via a Bond-Max autostaining system (Leica biosystems, Lincolnshire, IL). Stained kidneys were scanned at high resolution using a CRi Panoramic SCAN 40x Whole Slide Scanner (Perkin-Elmer).

Images were equalized in size and converted to .tif files using CaseViewer. Images were then imported into ImageJ, scale set for conversion between microns and pixels, and deconvoluted with the “H DAB” deconvolution option. A threshold of the resulting blue image was set at 215 to see how many pixels were negative for collagen I, and a threshold of the brown (IHC positive) image set at 185 to see how many pixels were positive for collagen I. Machine-staining allowed these kidneys to be compared with high reproducibility.

### Biochemistry analysis of serum for markers of kidney damage

At the time of euthanasia, blood was collected via submandibular bleed into protein low-bind tubes and allowed to coagulate for 2 hours on ice. Coagulated blood was then centrifuged at 10,000xg for 10 min, and serum collected. Serum was then diluted 4x in MilliQ water before being placed on deck on an Alfa Wassermann VetAxcel Blood Chemistry Analyzer. All tests requiring calibration were calibrated on the day of analysis and quality controls were run before analyzing samples. Serum tests were run according to kit instructions, and creatinine kinase concentration was normalized to calcium ion concentrations to account for sample hemolysis.

### Statistical analysis

Statistical analyses were performed using GraphPad Prism software, and *P* < 0.05 was considered statistically significant. Either Student’s t-test, 2-way ANOVA (with Fisher’s LSD post test), or 1-way ANOVA with Welch’s correction was used to compare groups.

## Acknowledgements

We thank the Human Tissue Resource Center of the University of Chicago for histology analysis. We thank the Integrated Light Microscopy Core of the University of Chicago for Imaging. We would also like to acknowledge guidance on fibrosis models from Dr. Anne I. Sperling. We thank the CTC Microsurgery Core at Northwestern University for performing the initial surgical procedures and University of Chicago Animal Research Center for help and assistance. We acknowledge Tera Lavoie from the Advanced Electron Microscopy Facility at the University of Chicago for technical assistance.

## Author’s contributions

Conceptualization: MJVW, MMR, EB, JAH

Methodology: MJVW, MMR, EB

Investigation: MJVW, MMR, EB, EY, AS, HS, ZJZ, LTG, SC, ATA

Visualization: MJVW, MMR, EB

Funding acquisition: JAH

Project administration: MJVW, JAH

Supervision: MJVW, JAH

Writing – original draft: MJVW, MMR, EB, LTG

Writing – review & editing: MJVW, MMR, EB, JAH

## Ethics approval

All the animal experiments performed in this work were approved by the Institutional Animal Care and Use Committee of the University of Chicago.

---

**Supplementary Figure 1.**
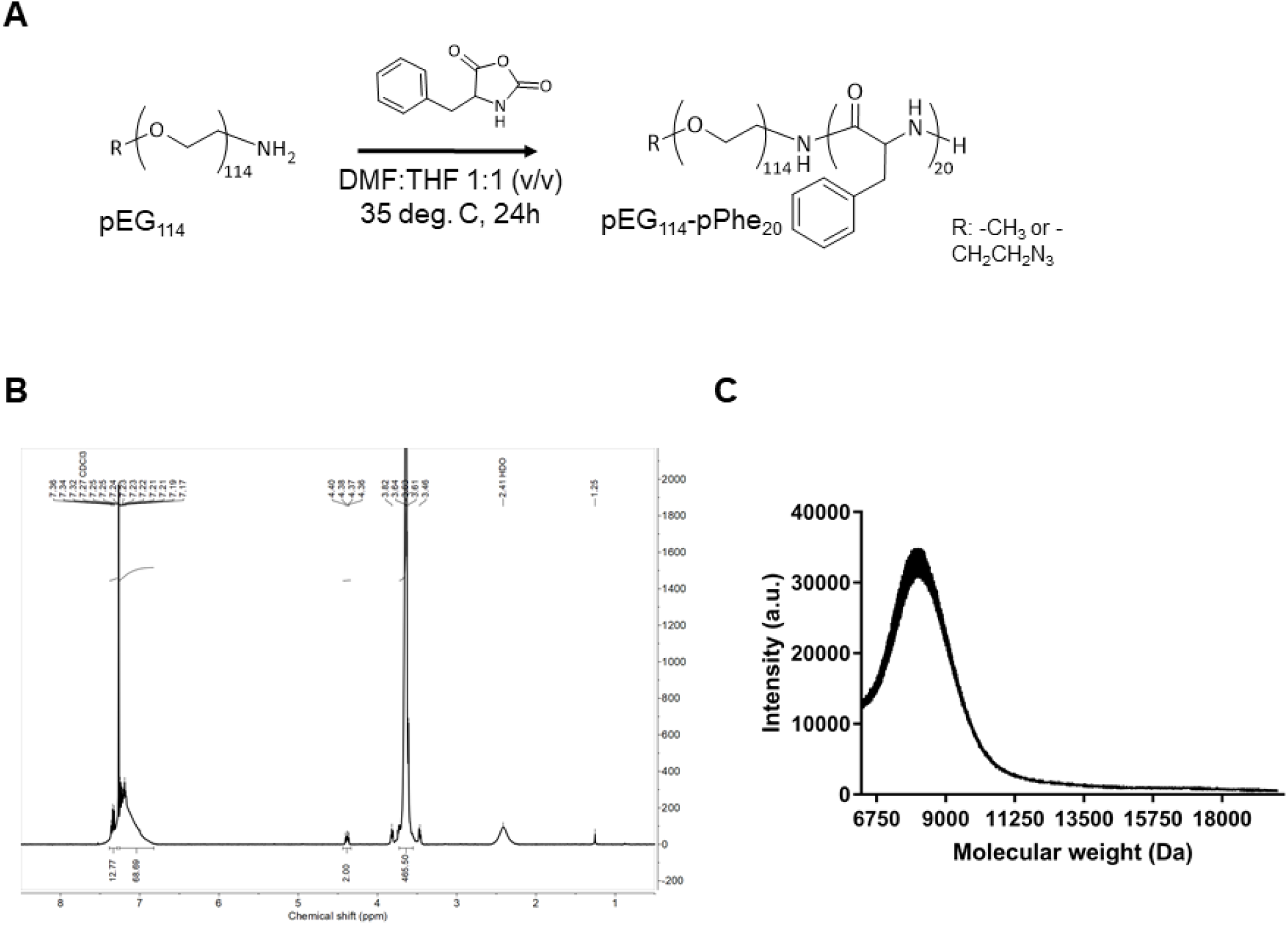
Synthesis and characterization of pEG_114_-pPhe_20_ block copolymer. **A**. Ring-opening polymerization of pEG_114_-pPhe_20_ from R-pEG_114_-NH_2_ and L,D-phenylalanine-N-carboxyanhydride (Phe-NCA), where R stands for either methoxy or azide terminal group. **B**. Proton NMR spectrum of N_3_-pEG_114_-pPhe_20_ block copolymer. Y axis represents intensity (a.u.) for the corresponding chemical shift values (parts per million, ppm). **C**. MALDI-TOF spectrum of R-pEG_114_-pPhe_20_ indicating the molecular weight of the block copolymer of about 8 kDa.

**Supplementary Figure 2.**
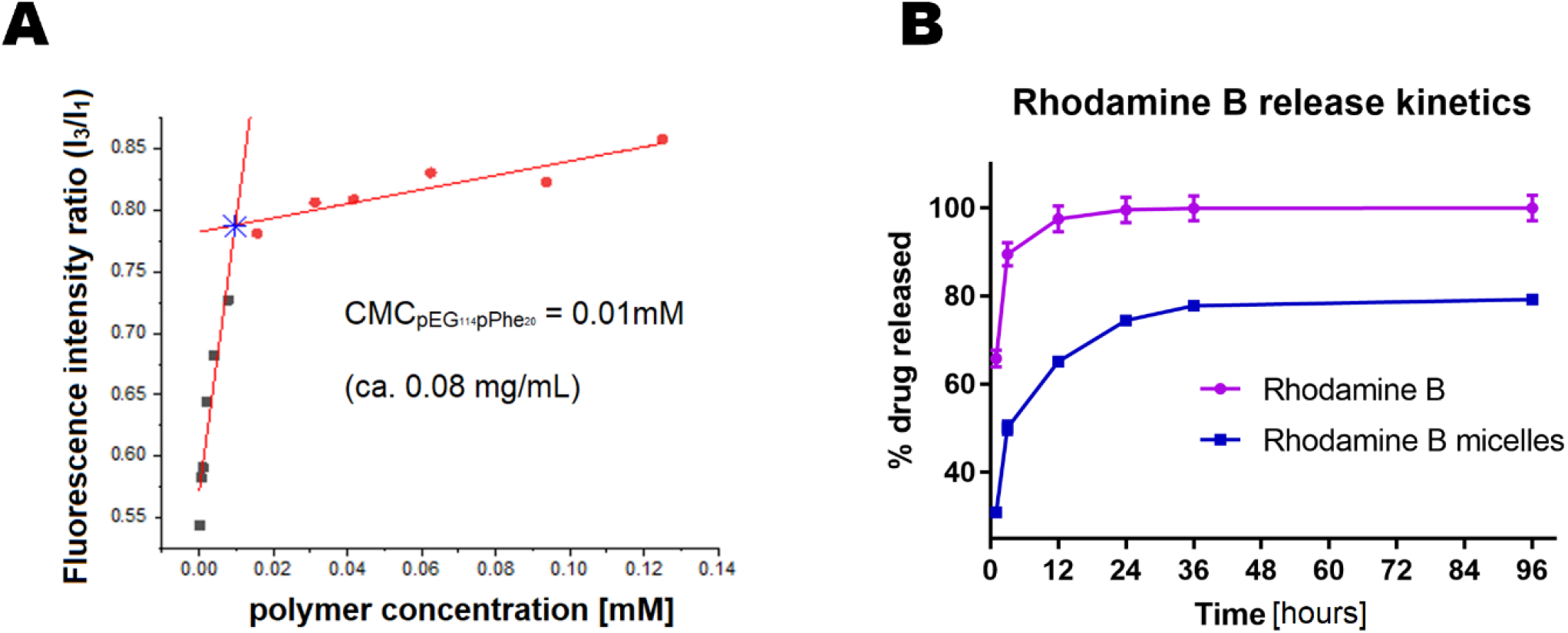
Characterization of pEG_114_-pPhe_20_ micelles as drug delivery system. **(A)** Determination of critical micelle concentration (CMC) of pEG_114_-pPhe_20_ micelles using pyrene fluorescence technique. Ratio of the fluorescence intensity of band 3 (I_3_) to band 2 (I_1_) of pyrene was plotted against polymer concentration in measured solution. CMC was determined by conducting linear regression calculations (red lines), where CMC is showed as a blue star. (B) Drug release profile of Rhodamine B-encapsulated pEG_114_-pPhe_20_ micelles (Rhodamine B micelles) compared to drug release profile of free Rhodamine B. Drug-micelles or free drug was dialyzed against PBS and the fluorescence at ex/em 550/600 of eluted drug was measured and quantified as % drug release.

**Supplementary Figure 3.**
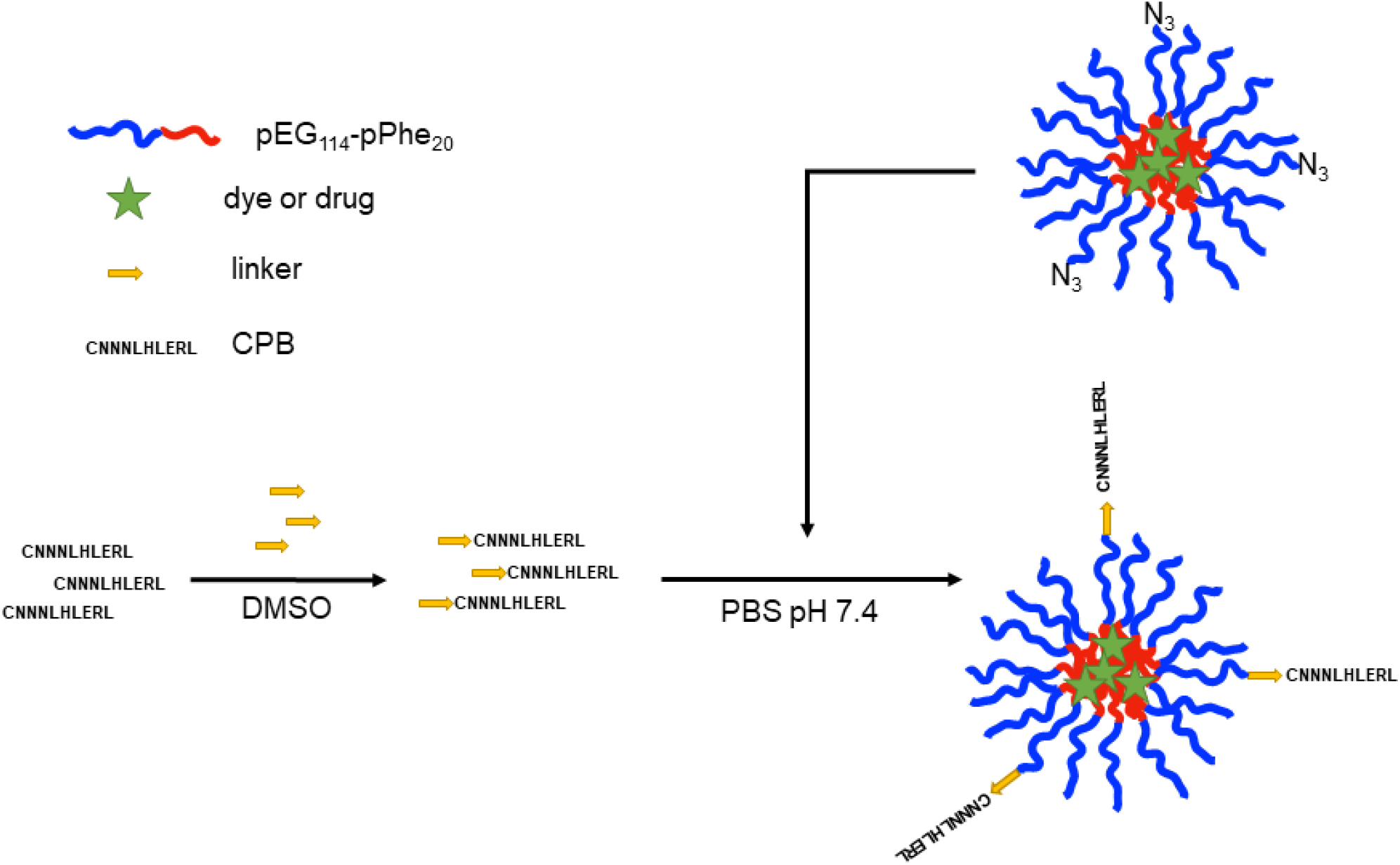
Schematic describing a strategy to formulate peptide-conjugated pEG_114_-pPhe_20_ micelles used in this study. DMF: dimethylformamide, THF: tetrahydrofuran, PBS: phosphate-buffered saline, N_3_:azide, DMSO: dimethyl sulfoxide, CBP: collagen-binding peptide.

**Supplementary Figure 4.**
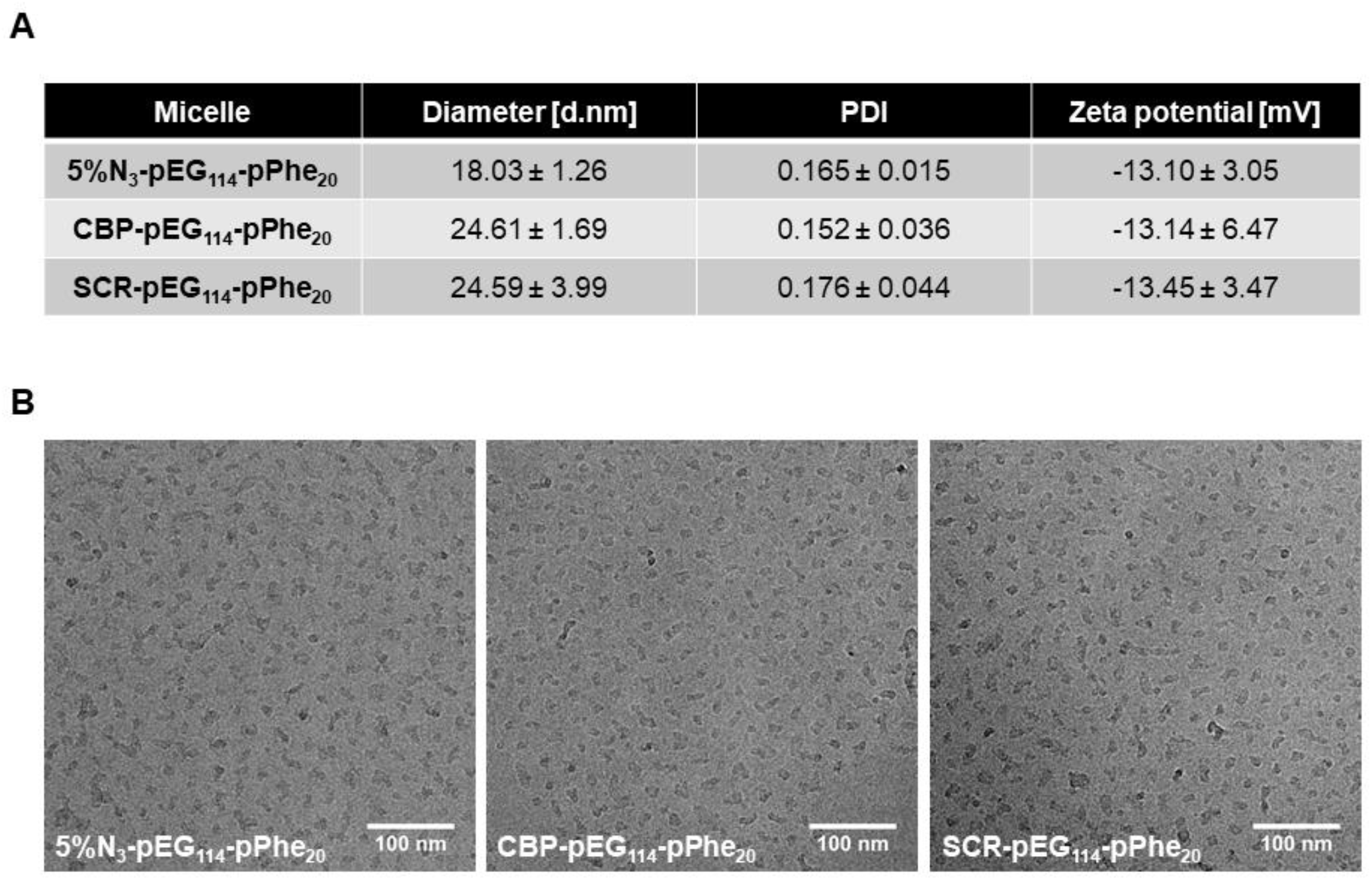
Characterization of peptide-conjugated micelles. **A**. Nanoparticle diameter (in nm), polydispersity index (PDI, a.u.) and Zeta potential (mV) of 5%N_3_-pEG_114_-pPhe_20_ (unconjugated), CBP-pEG_114_-pPhe_20_ and SCR-pEG_114_-pPhe_20_ micelles, determined by Dynamic Light Scattering measurements. B. Representative cryo-EM images of 5%N_3_-pEG_114_-pPhe_20_ (unconjugated), CBP-pEG_114_-pPhe_20_ and SCR-pEG_114_-pPhe_20_ micelles. Scale bar: 100 nm.

**Supplementary Figure 5.**
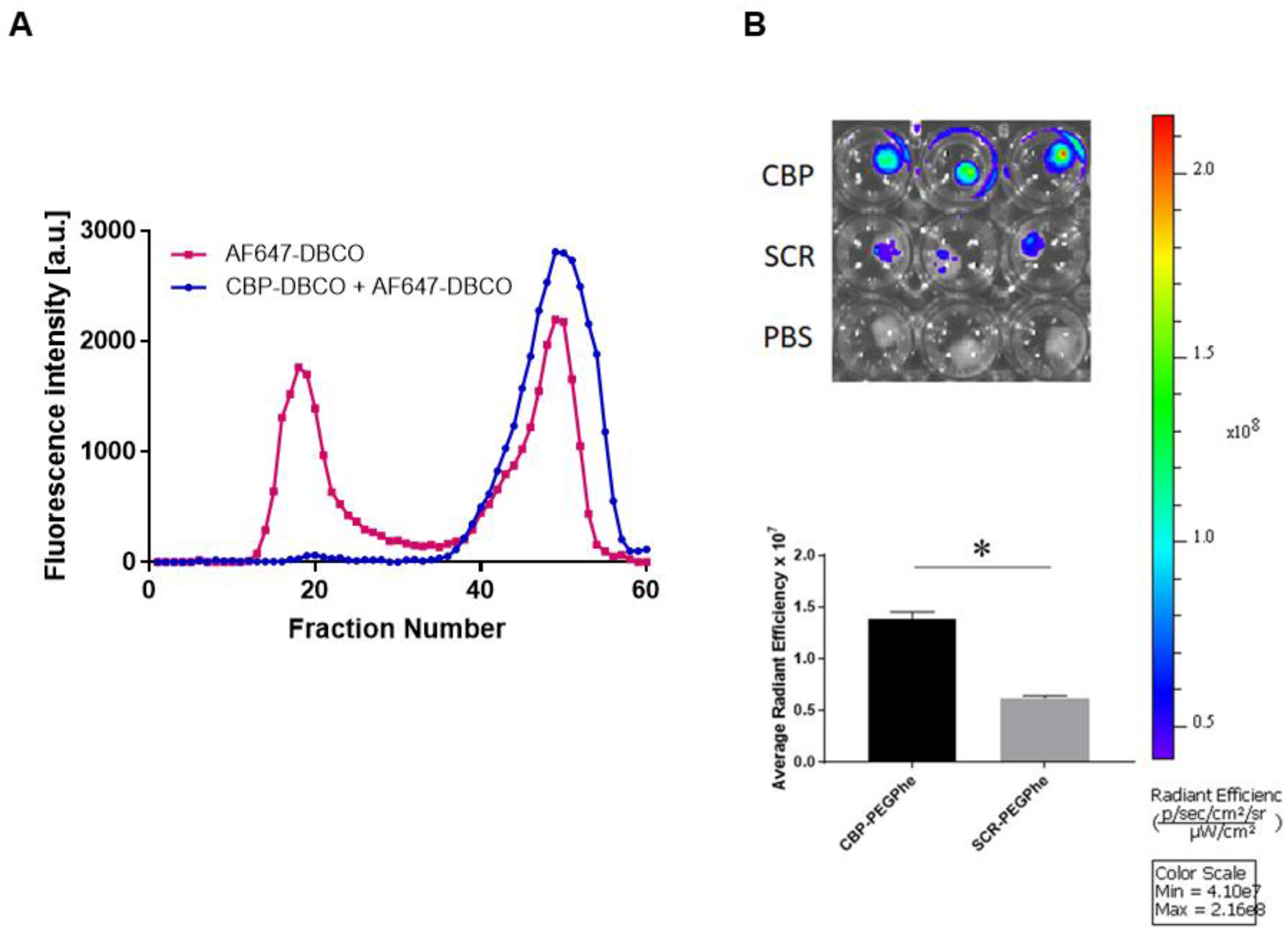
CBP-conjugated micelles bind to collagen type I. A. SEC elution profile of azide-modified micelles (pink) or CBP-micelles (blue) conjugated to AF647-DBCO dye. Fractions 15-25 correspond to micelles. Fractions 40-60 correspond to unreacted dye. Fluorescence detection was performed via fluorescence plate reader at 650/665. One fraction corresponds to ca. 0.25 mL of elution volume. B. Fluorescence intensity of collagen type I sponges incubated with AF647-conjugated micelles conjugated to either CBP or SCR peptide. The signal was measured after washing the sponges with PBS by IVIS imaging system. The sponges soaked in PBS were used as background. Top image depicts fluorescence intensity (in radiant efficiency, right side) of the collagen sponges. Botton graph is a numerical representation of the top image and shows average radiant efficiency of CBD-micelle-and SCR-micelle-incubated collagen sponges. Comparison was made by Student’s t-test with * at p<0.05

**Supplementary Figure 6.**
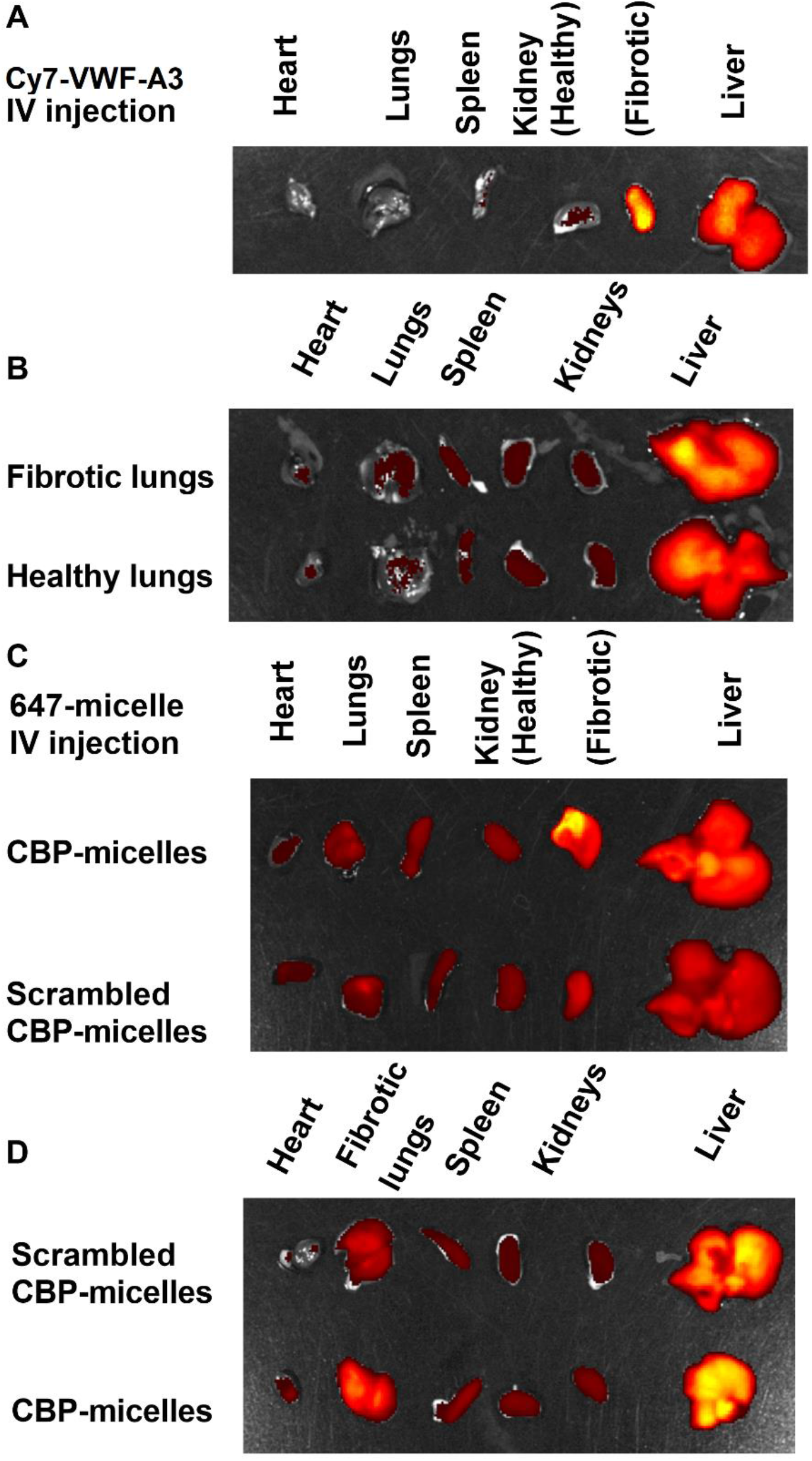
Fluorescently labeled VWF-A3, scrambled CBP-micelles, and CBP-micelles target to both kidney and lung fibrosis. Mice with fibrotic kidneys, fibrotic lungs, or healthy lungs were injected 1 week after insult (or sham) with fluorescently labeled VWF-A3, scrambled CBP-micelles, and CBP-micelles. Heart, lung, spleen, kidneys, and liver were harvested 24 hours after injection for fibrotic kidney experiments or 48 hours for fibrotic lung experiments. Fluorescence intensity was measured via IVIS. Cy7-VWF-A3 was found in higher concentrations in both (A) fibrotic kidneys and (B) lungs. 647-scrambled-CBP-micelles and, to a greater degree, 647-CBP-micelles were found in higher concentrations in both (C) fibrotic kidneys and (D) lungs. Measurements can be found in Figure 1.

**Supplementary Figure 7.**
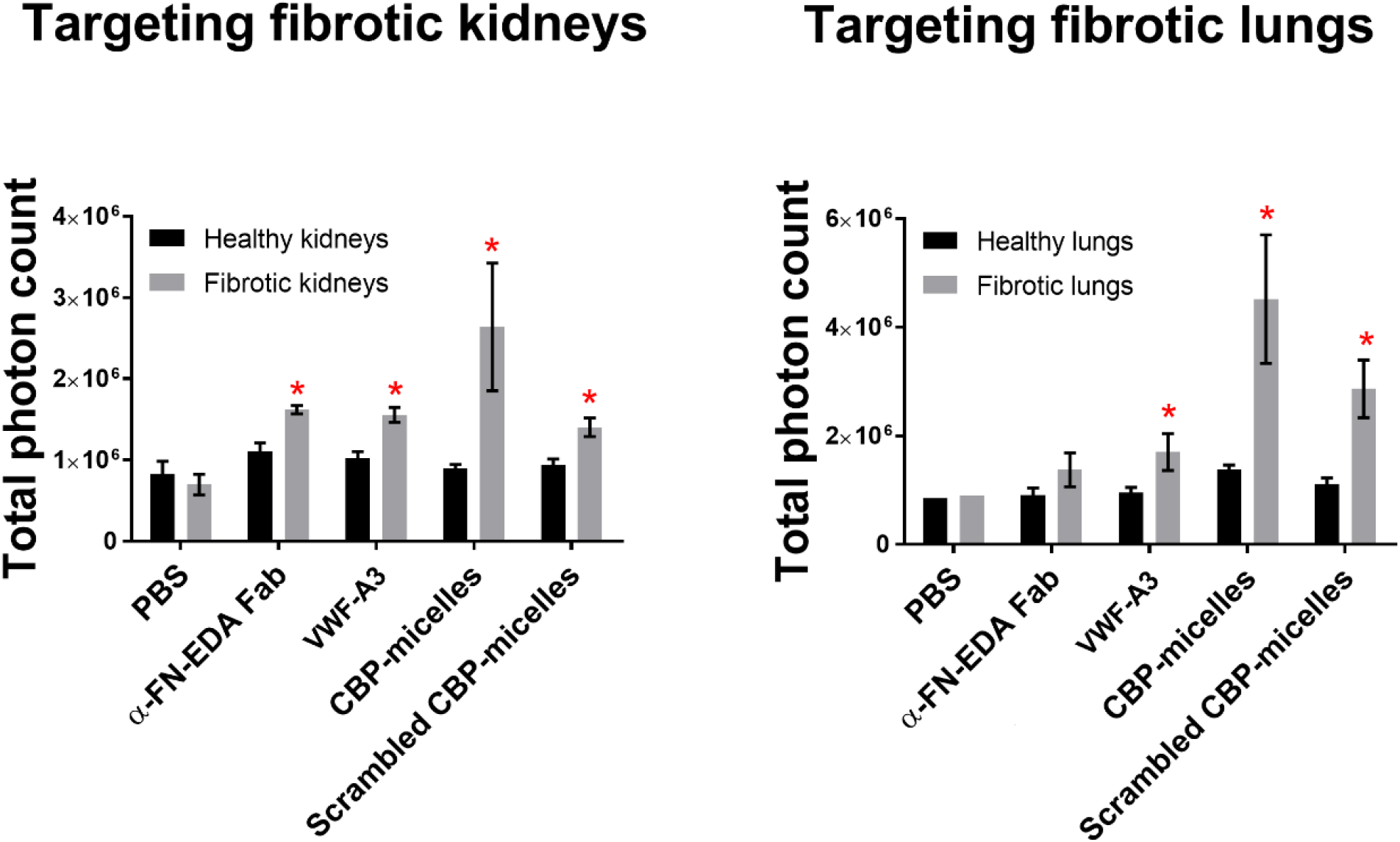
Fluorescently labeled VWF-A3, scrambled CBP-micelles, and CBP-micelles target to both kidney and lung fibrosis. Data from Figure 1 presented as a total photon count that is not normalized to total fluorescent intensity.

**Supplementary Figure 8:**
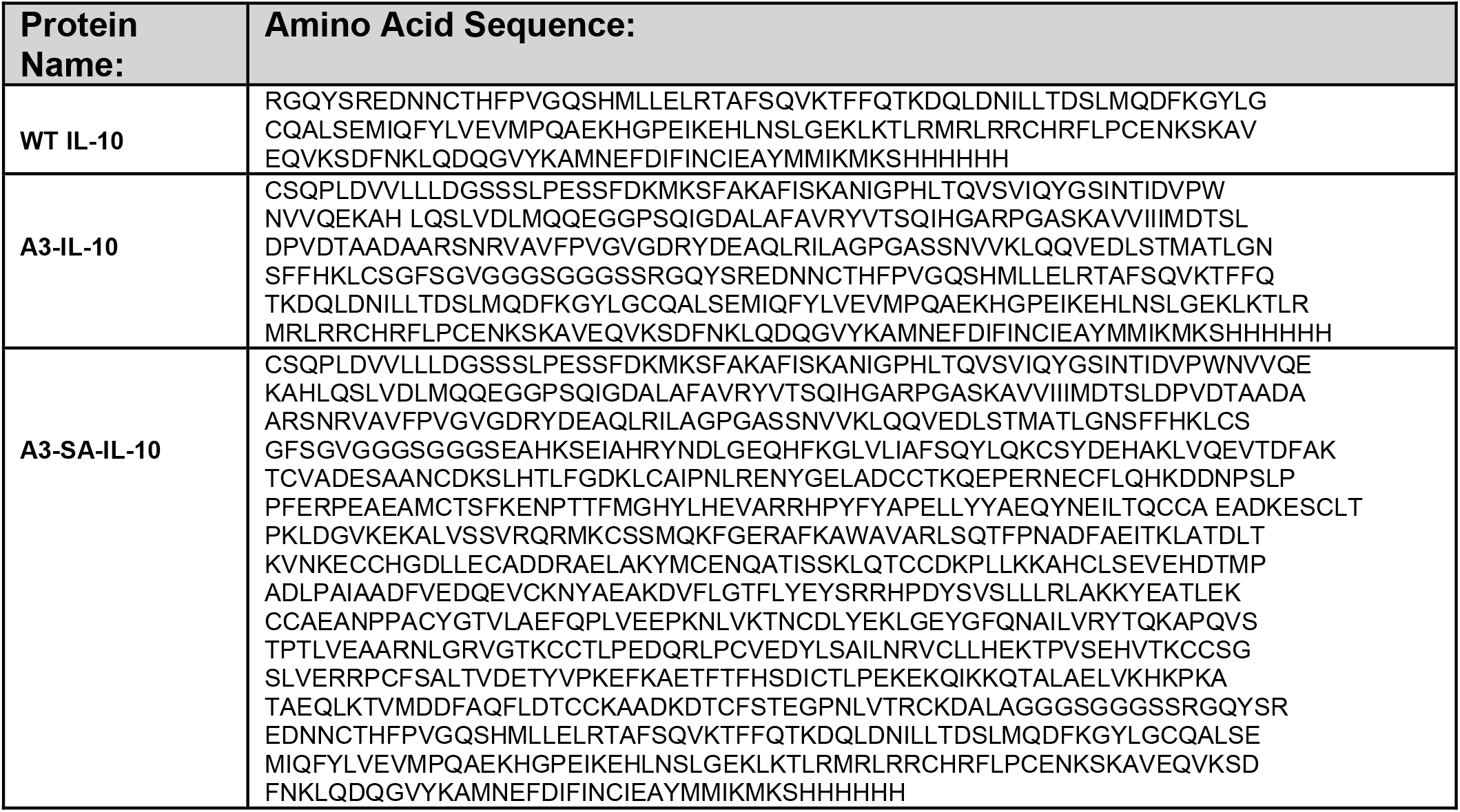
Amino acid sequences of unmodified mouse IL-10, A3-IL-10 and A3-SA-IL-10.

**Supplementary Figure 9.**
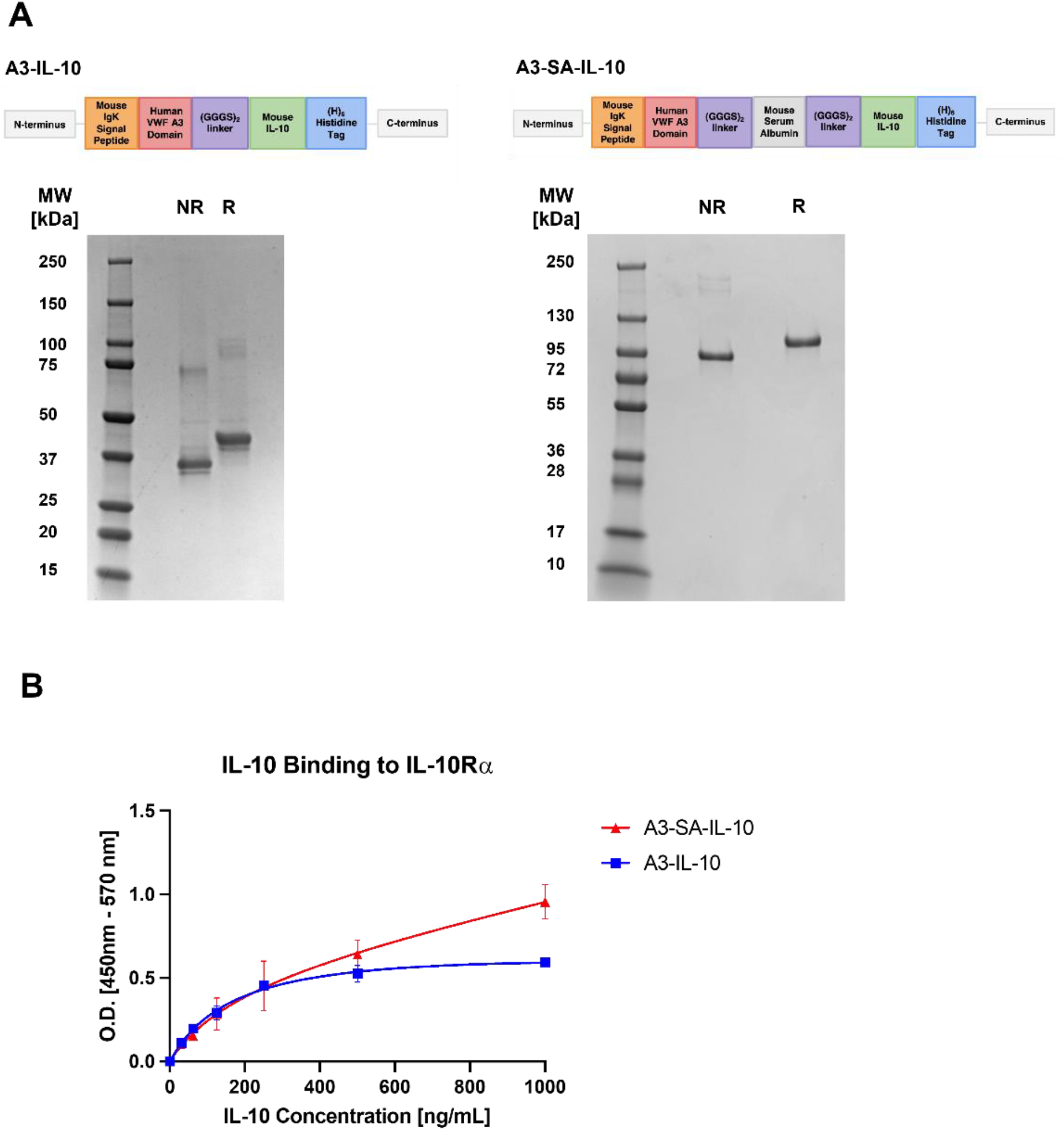
A3-IL-10 and A3-SA-IL-10 bind to IL-10 receptor. **(A)** SDS-PAGE image of A3-IL-10 (left) and A3-SA-IL-10 (right). NR indicates non reducing conditions, R reducing conditions, (B) ELISA-based affinity measurements of A3-SA-IL-10 and A3-IL-10 binding to IL-10Ra.

**Supplementary Figure 10.**
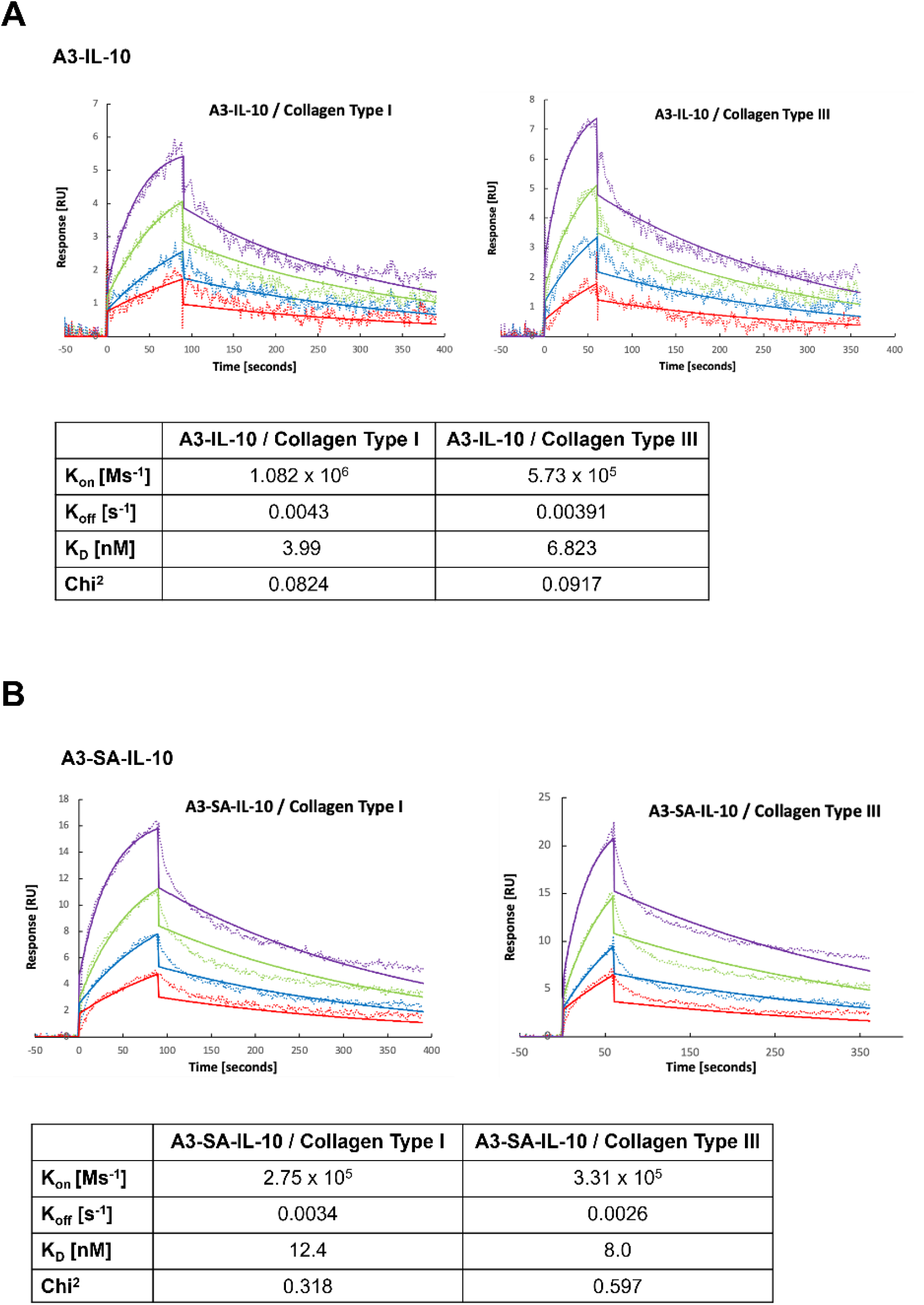
A3-IL-10 and A3-SA-IL-10 bind collagen type I and III. SPR analysis of (A) A3-SA-IL-10 and (B) A3-IL-10 binding to collagens I and III.

**Supplementary Figure 11.**
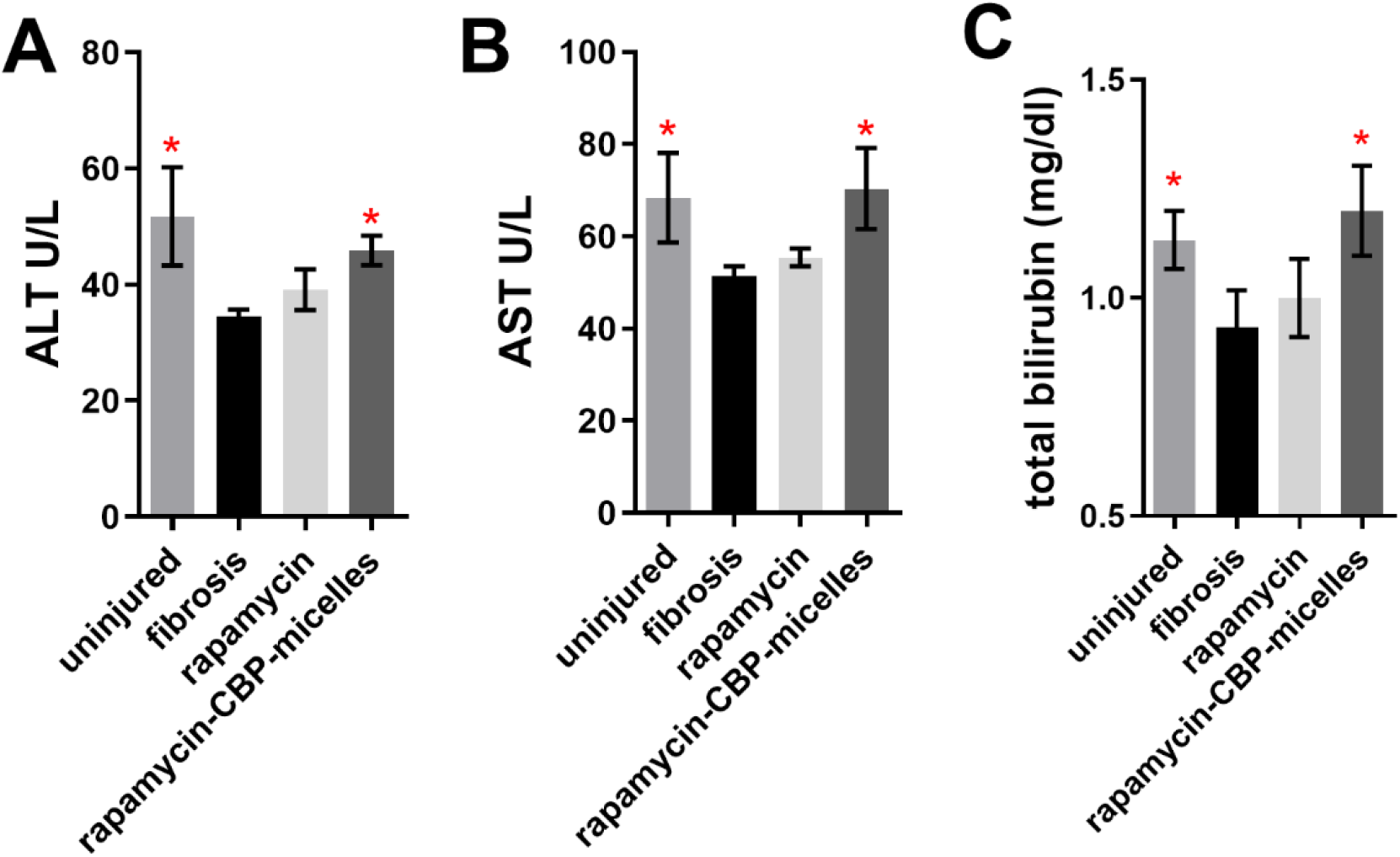
In mice with a fibrotic left kidney, rapamycin-CBP-micelle treatment restores ALT, AST, and bilirubin concentrations to levels comparable to healthy mice. Blood was taken from mice directly before sacrifice at 2 weeks post UUO ligation. Blood was also taken from age-matched mice with uninjured kidneys. Concentrations in serum of (A) ALT, (B) AST, (C) bilirubin. * = statistical significance of P < 0.05, < 0.01, or < 0.001, significance vs fibrosis, one way ANOVA, no post test.

**Supplementary Figure 12.**
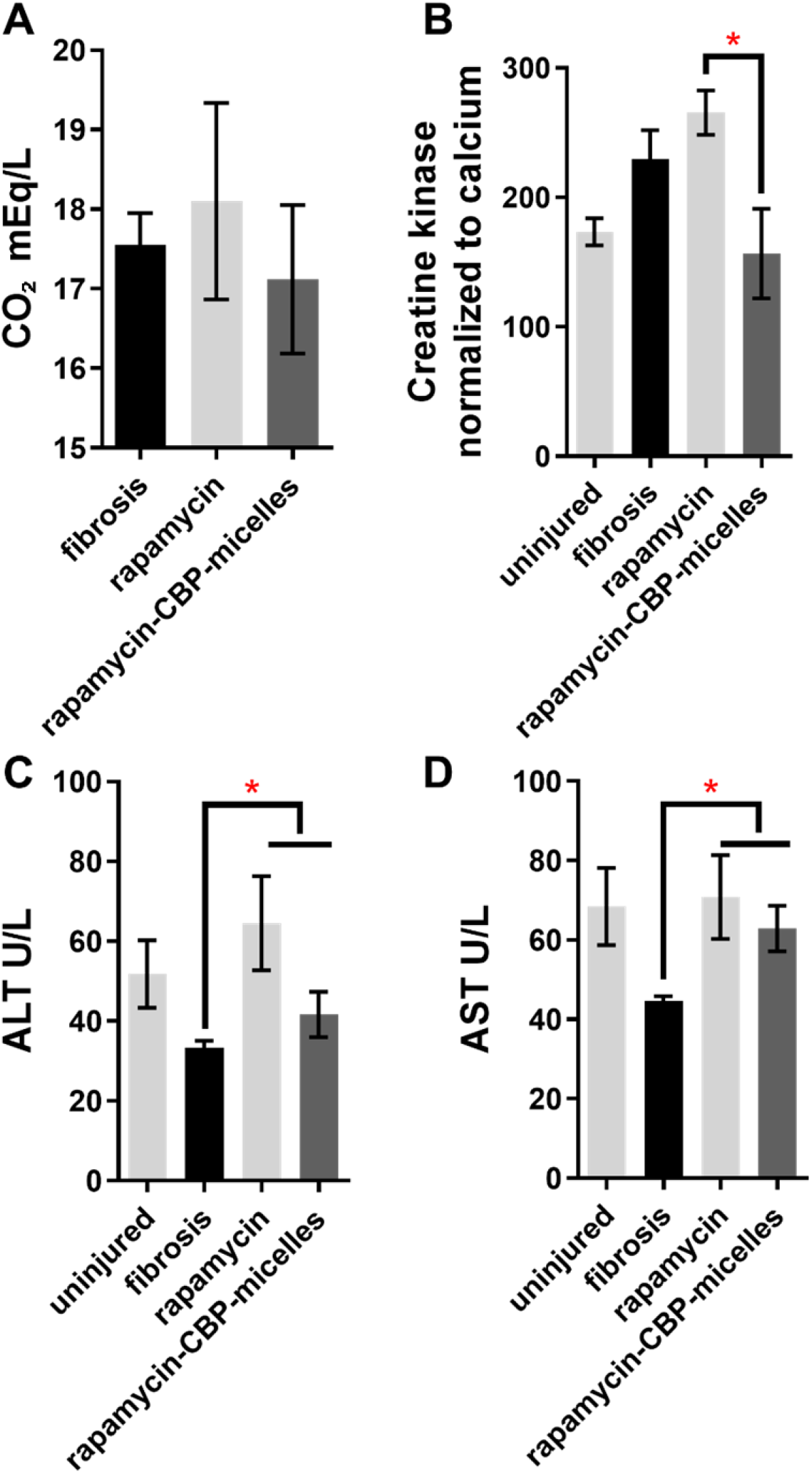
In mice with fibrotic lungs, rapamycin-CBP-micelles treatment restores creatine kinase, ALT, and AST concentrations to levels comparable to healthy mice. Blood was taken from mice directly before sacrifice at 3 weeks post bleomycin insult. Blood was also taken from age-matched mice with uninjured lungs. Concentrations in serum of (A) CO_2_, (B) creatine kinase, (C) ALT, and (D) AST. * = statistical significance of P < 0.05, significance vs fibrosis, one way ANOVA, no post test. Comparison in (B) is Student’s t-test

**Supplemental Figure 13.**
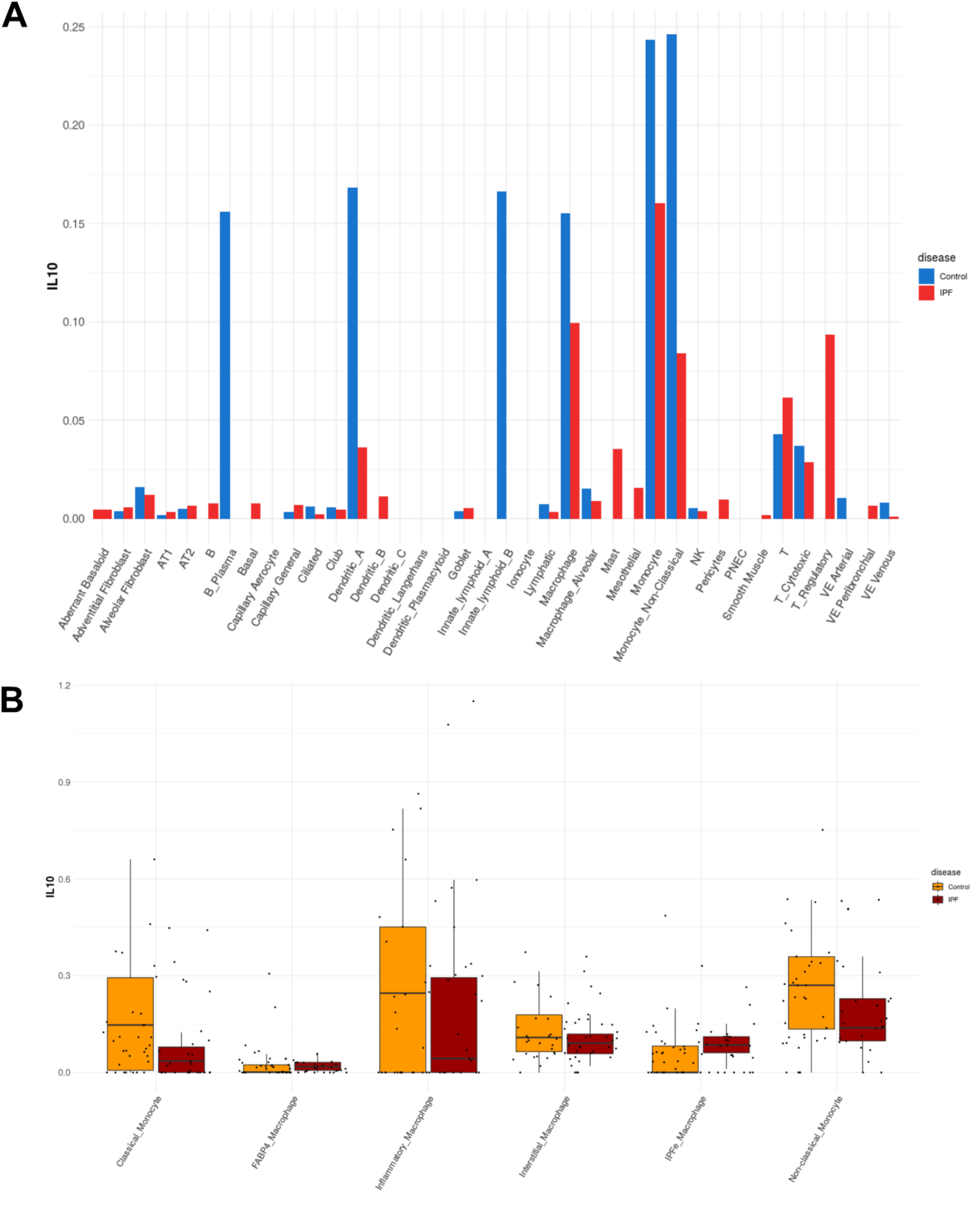
IL10 is downregulated among many cell types found in IPF. (A) IL10 transcript frequency in cells found in IPF. (B) IL10 transcript frequency by individual patient for selected immune cell subtypes. All data and figures from the IPF Atlas, Kraminski/Rosas dataset.

**Supplemental Figure 14.**
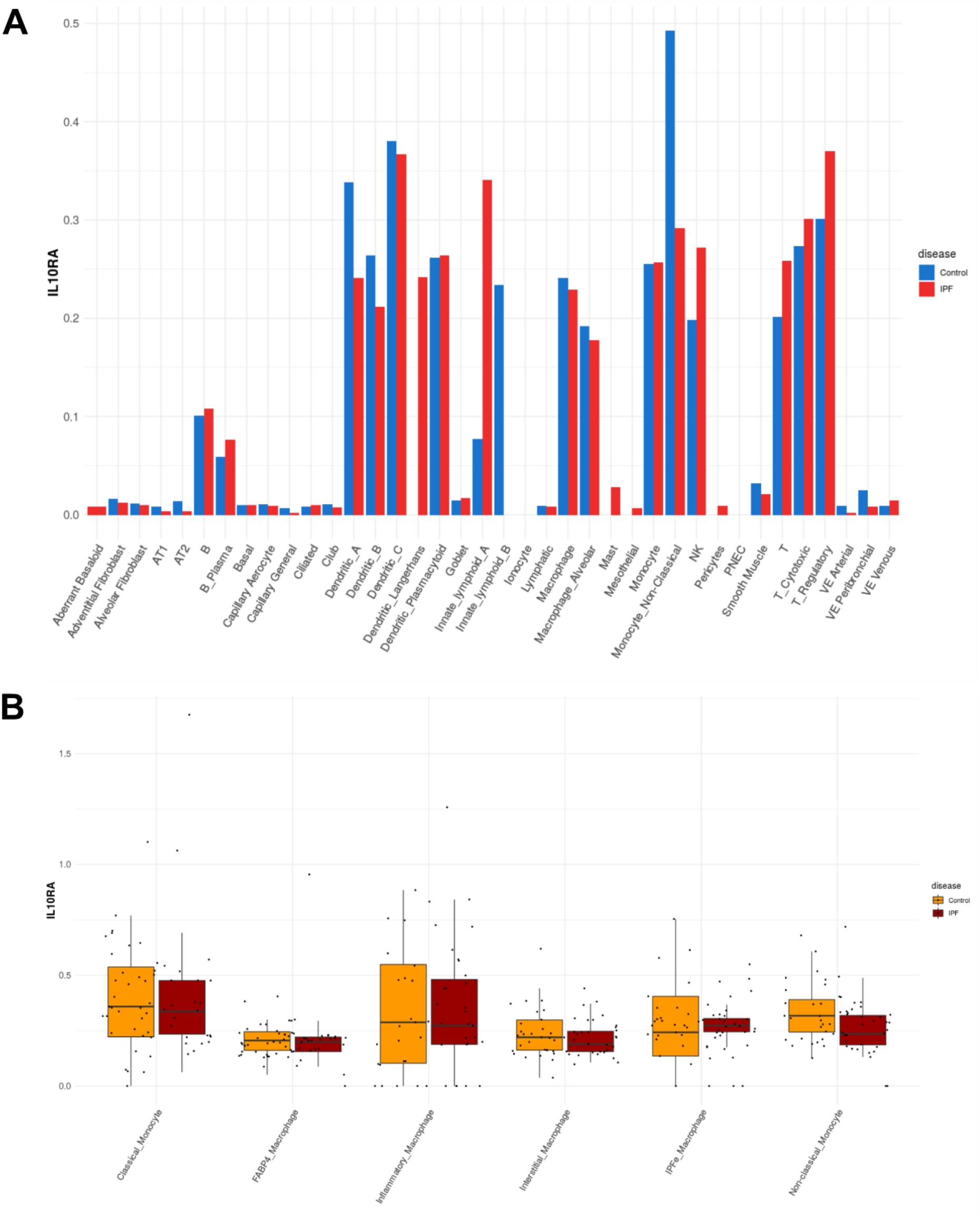
IL10 receptor is downregulated among many cell types found in IPF. (A) IL10 receptor transcript frequency in cells found in IPF. (B) IL10 receptor transcript frequency by individual patient for selected immune cell subtypes. All data and figures from the IPF Atlas, Kraminski/Rosas dataset.

**Supplemental Figure 15.**
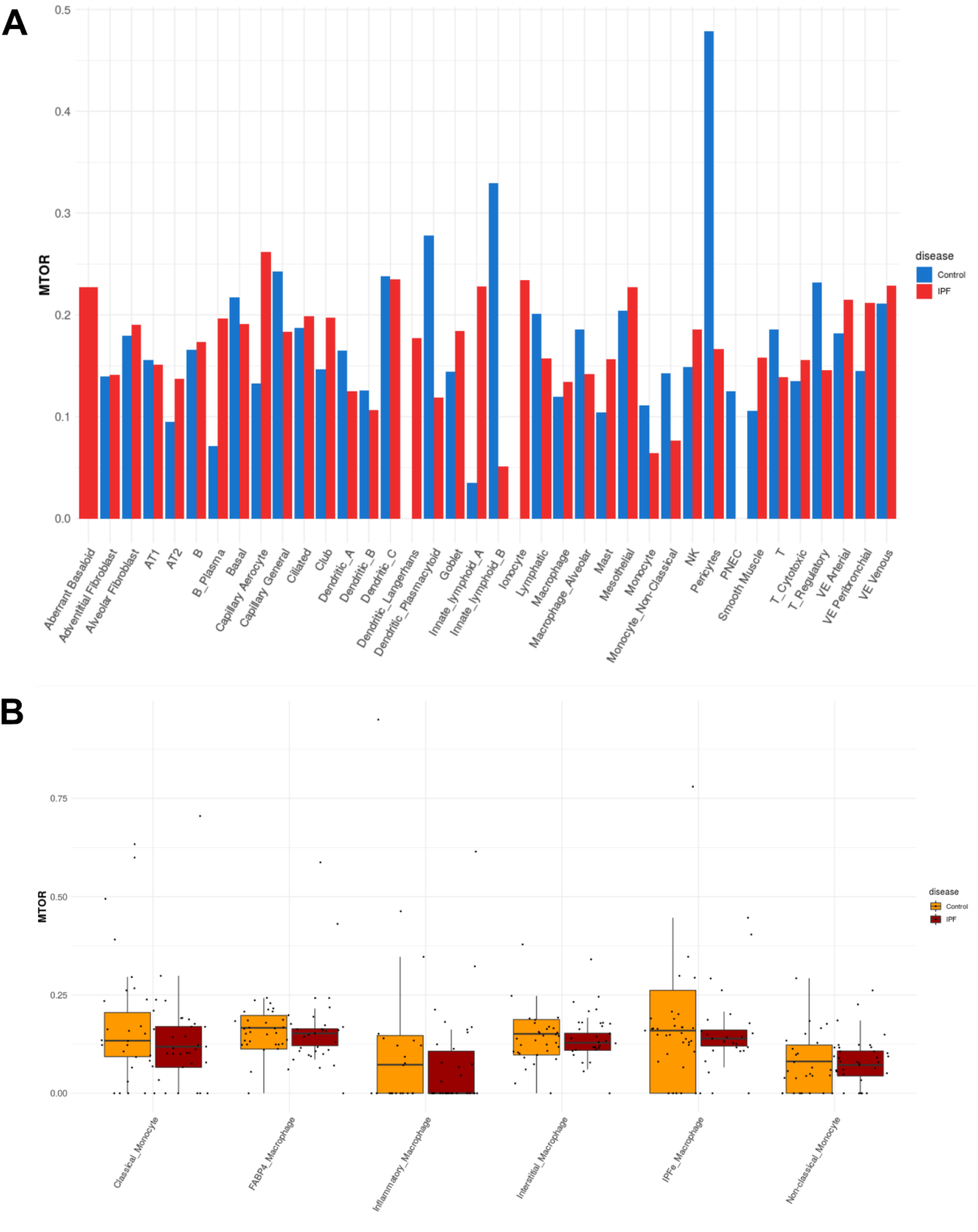
mTOR receptor is downregulated among many cell types found in IPF. (A) mTOR receptor transcript frequency in cells found in IPF. (B) mTOR receptor transcript frequency by individual patient for selected immune cell subtypes. All data and figures from the IPF Atlas, Kraminski/Rosas dataset.

